# HYphlow: bringing end-to-end flow to HyPhy selection analyses, from taxon reconciliation to trait-based foreground assignment

**DOI:** 10.64898/2026.10.30.763228

**Authors:** Hyejung Kwon, Taegan JM Perez, Eren Yigit, Ryan K. Schott

## Abstract

Phylogenetic analyses of molecular evolution often require repeated manual preparation to reconcile taxa, annotate trees, execute models, and summarize results. These steps become increasingly time-consuming and difficult to standardize as the number of taxa, genes, and analytical hypotheses increases. We present HYphlow, an open-source graphical workflow developed in Python that integrates data preparation, trait-based foreground branch assignment, HyPhy execution, and structured result extraction within a single interface. HYphlow reconciles taxon labels across alignments, trees, and trait data; reconstructs ancestral states using parsimony and maximum likelihood; generates alternative foreground annotations; executes HyPhy analyses locally or through portable scripts for high-performance computing; and records analytical decisions and settings throughout the workflow. We evaluated HYphlow using simulations and two published datasets comprising cryptochrome analyses in pigeons and doves and visual opsin analyses in skinks. A name-similarity threshold of 80 reliably recovered simulated spelling variants, while consensus foreground assignment increased precision relative to parsimony or maximum likelihood alone (0.928 *vs* 0.891 and 0.907, respectively), with a corresponding reduction in recall (0.875 *vs* 0.921 and 0.918). In the empirical datasets, HYphlow reconciled taxon labels, reconstructed or reproduced foreground hypotheses derived using different approaches, and generated selection results consistent with the principal conclusions of the original studies. HYphlow therefore provides a reproducible framework for carrying comparative molecular evolutionary analyses from independently prepared sequence, phylogenetic, and trait data through branch assignment, model execution, and results summary while retaining researcher control over analytical decisions. HYphlow is freely available under the MIT licence from GitHub at: https://github.com/Schott-Lab/HYphlow.

## Introduction

To test whether a trait affects selection on a gene, researchers identify the branches associated with that trait. Codon models use the ratio of nonsynonymous to synonymous substitution rates (dN/dS or *ω*) to estimate selective pressure (Nei and Gojobori 1986; Goldman and Yang 1994; Muse and Gaut 1994; Yang and Nielsen 2000). This ratio can vary across sites and evolutionary lineages (Yang et al. 2000; Kosakovsky Pond et al. 2011). Branch-site models account for variation in *ω* across sites and branches (Yang 1998; Yang and Nielsen 2002; Zhang et al. 2005). Branches selected for testing are called foreground branches.

HyPhy (H**y**pothesis Testing using Phylogenies) provides methods for testing selection on specific branches (Kosakovsky Pond et al. 2020). For example, BUSTED (Branch-Site Unrestricted Statistical Test for Episodic Diversification) tests for episodic diversifying selection on specified branches (Murrell et al. 2015). RELAX compares the strength of selection between foreground (test) and reference branches (Wertheim et al. 2015). When studying a trait such as habitat or diet, researchers can define foreground branches based on which lineages are inferred to carry the trait. The selected branches determine which lineages are included in the test.

Defining these branches requires trait data, while HyPhy uses a codon alignment and a phylogenetic tree with labelled branches for the selection test. These inputs may come from different sources, and foreground branches may need to be assigned separately for each gene.

Linking these inputs depends on taxon labels. Sequence labels, tree tip labels and species names in trait tables may use different formats or abbreviations for the same taxon. Taxonomic issues such as misspellings, synonyms and revised names can make matching more difficult (Boyle et al. 2013; Remsen 2016; Grenié et al. 2023). For example, sequences from GenBank may use names recorded in NCBI (National Center for Biotechnology Information) Taxonomy (Schoch et al. 2020), whereas trait data compiled from other databases may follow another taxonomy. These naming issues have been documented in vertebrate occurrence records. Among 1,000 name combinations published through VertNet, 991 could be assessed and 532 contained at least one naming issue. Of the 1,000, 33 remained unresolved after manual review (Zermoglio et al. 2016). Matching sequence labels, tree tips and trait records therefore involves decisions about which names refer to the same taxon. However, the alignment, tree and trait table do not record how these decisions were made.

Trait data describe the taxa at the tips of a tree, whereas foreground branches can also include internal branches. The foreground status of internal branches therefore depends on ancestral states inferred by ASR (ancestral state reconstruction) (Pagel 1999). Ancestral states can be reconstructed manually, using custom scripts, or using parsimony and likelihood methods. Different methods can assign different states to the same node, and some nodes may remain ambiguous (Cunningham et al. 1998; Joy et al. 2016). ASR estimates states at nodes, while foreground labels are assigned to branches. When a trait changes along a branch or the ancestral state is ambiguous, whether the branch belongs to the foreground depends on a labeling rule.

When alignments differ in species sampling, the pruned tree and the reconstructed states can also differ among genes, and gene trees can differ in topology from the species tree (Avise and Robinson 2008; Mendes and Hahn 2016). The same trait hypothesis can therefore be tested on different sets of branches, and each set poses a different hypothesis to the selection analysis. When branch sets differ among genes, the labelled trees alone cannot show whether the differences arose from the trees, the trait data or the labelling rule.

Comparative studies repeat these analyses across genes. Each gene can be tested with several methods and trait hypotheses, so the number of analyses grows with every gene, method and branch set added. Large sets of analyses can be run in parallel on local computers or computing clusters. HyPhy writes a separate result file for each analysis, which records the input tree, the tested branches and the fitted models, but not how taxon names were matched or why each internal branch was assigned to the foreground. Model fitting adds a further source of variation. Some HyPhy methods start parameter estimation from randomly chosen values (Castoe et al. 2008), and repeated analyses of the same data can produce different estimates (Selberg et al. 2025). In maximum likelihood phylogenetic inference, random seeds and the number of processor threads can affect reproducibility but are rarely reported (Shen et al. 2020). When taxon matches, branch assignments and random seeds are kept outside the result files, differences between analyses can be difficult to trace.

Preparing these inputs and running the selection analyses can involve several programs, each supporting part of the process. The names can be matched against reference taxonomies with the TNRS (Taxonomic Name Resolution Service; Boyle et al. 2013) and taxize (Chamberlain and Szöcs 2013). For codon-based selection analyses with PAML (Phylogenetic Analysis by Maximum Likelihood; Yang 2007), graphical and command-line interfaces such as PAMLX (Xu and Yang 2013), EasyCodeML (Gao et al. 2019), BlastPhyMe (Schott et al. 2019), and ete-evol (Huerta-Cepas et al. 2016) configure and run codeml analyses, with varying support for data preparation. HyPhy provides methods for testing selection across sites and branches, including tests for relaxed selection, within a single program (Wertheim et al. 2015; Kosakovsky Pond et al. 2020). Tools built around HyPhy support different stages of analysis, including running selection tests from the command line, a web browser or a workflow system, labelling branches, and displaying results (Table 1). Some tools support individual steps, while others combine several steps into a single workflow. Scripts can also connect separate programs and record the steps and settings used. However, recording researcher decisions involves more than saving the commands and settings used. Connecting these decisions to individual selection results requires additional steps.

**Table 1.** Comparison of Pros and Cons of Different Approaches to Trait-Based Selection Analyses.

| Approach | Pros | Cons |
|---|---|---|
| Manual branch selection | Researchers assess each branch directly. | Repeated for each gene and tree; the basis for each label is not stored with the tree. |
| Scripted ancestral state reconstruction | Flexible choice of reconstruction models; steps and settings are stored in code. | Requires programming; decisions made between steps are stored separately from selection results. |
| Command line with labelling scripts | Rule-based labelling of internal branches; runs locally. | Name matching and per-gene tree preparation are performed separately. |
| Web server | No local installation; interactive branch selection. | Branches are selected separately for each analysis. |
| End-to-end pipeline | Automates sequence alignment, tree inference, selection analyses and summary. | Intermediate steps follow fixed settings, with limited points for researcher review. |

Here, we present HYphlow, a graphical workflow for trait-based selection analyses with HyPhy, in which each selection result is linked to the decisions behind it. It builds on HyPhy’s selection tests and on established methods for taxon name matching and ASR. Steps that follow fixed rules are automated, while decisions that require researcher judgement remain under researcher control. Both automated steps and researcher decisions are recorded. The workflow consists of four stages: data preparation, tree annotation, HyPhy execution and results summary.

In data preparation, taxon labels are reconciled across the trait table, alignments and trees. Exact matches are accepted automatically, while approximate matches are proposed for the researcher to accept or decline. A shared species tree can also be pruned to the species in each alignment. In tree annotation, ancestral states are reconstructed for each gene, and foreground branches are assigned using the reconstruction method and labelling rule selected by the researcher. Inferred ancestral states and branch assignments are reported. In HyPhy execution, analyses across genes, methods and branch sets are run in parallel using a generated script that the researcher can review and edit. The random seed and number of processor threads used for each analysis are recorded. In results summary, results are collected into a single table linking each analysis to its gene, branch set and settings.

We evaluate each stage of HYphlow against known or published results. Using simulated datasets with known taxon identities and ancestral states, we measure how often proposed name corrections identify the correct taxon, how often incorrect suggestions occur, and how accurately foreground assignments reflect the simulated trait history. We then test whether HyPhy analyses run through HYphlow produce the same results as direct HyPhy analyses using the same random seed. Finally, we apply HYphlow to two published datasets, from their original taxon labels through selection analyses. In the cryptochrome study (Sweet et al. 2026), foreground branches were identified using likelihood-based ASR and labelled only where the trait changed. In the visual opsin study of *Lerista* and *Ctenotus* skinks (Ford et al. 2026), foreground branches were selected manually and included both terminal and internal branches within each group. For each dataset, we compared foreground branch sets produced by different reconstruction methods and labelling rules, including those used in the original study.

## Methods

### Implementation, installation and availability

HYphlow was developed and tested with Python v3.12.12, and its graphical interface uses PyQt5 v5.15.11 (Riverbank Computing Limited 2024). Taxon names were checked with taxopy v0.14.0 (Camargo 2024), labels are compared with RapidFuzz v3.14.3 (Bachmann 2025), and trees were handled with ETE3 (Environment for Tree Exploration) v3.1.3 (Huerta-Cepas et al. 2016).

Selection analyses are run with HyPhy, and v2.5.94 was used in this study (Kosakovsky Pond et al. 2020). HYphlow runs on macOS and Linux, and on Windows through the WSL (Windows Subsystem for Linux). It can be installed with a single command from Bioconda (`conda install -c conda -forge -c bioconda hyphlow`), which installs HyPhy as a dependency if it is not already installed in the environment. Alternatively, HYphlow can be installed from the Python Package Index (PyPI) using `pip install hyphlow`, with HyPhy installed separately. The source code is available on Github under the MIT License, and the version used in this study was v1.1.0 (Fig. 1).

**Figure 1.**
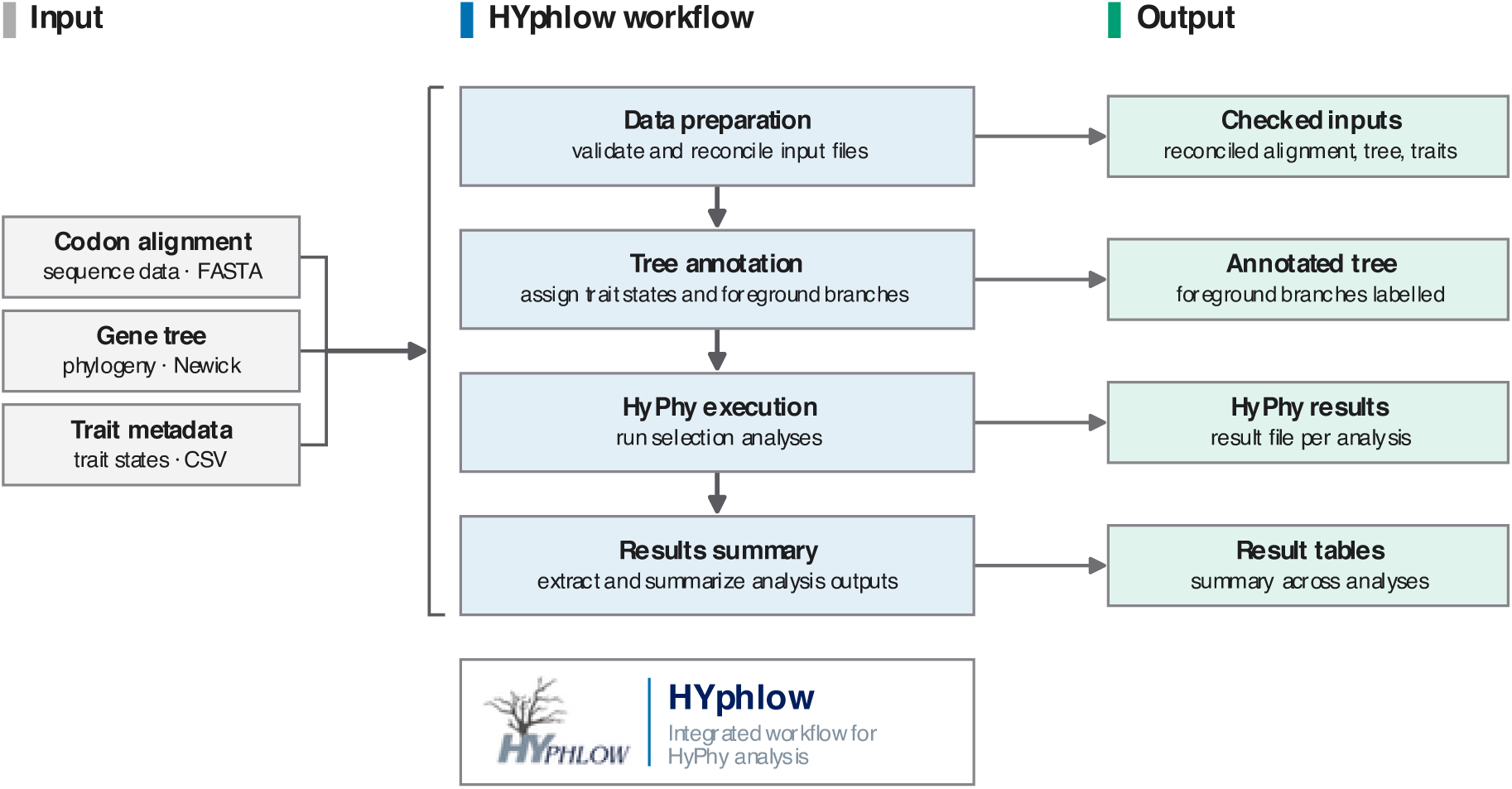
Overview of the HYphlow workflow. Codon alignments (FASTA), phylogenetic tree (Newick) and a trait table (CSV) (left) pass through four stages (centre), with outputs from each stage shown on the right. In data preparation, taxon labels are compared across the three inputs to identify naming differences. Exact matches are accepted automatically, and approximate matches are suggested for the researcher to accept or decline. In tree annotation, ancestral states are reconstructed using maximum parsimony and maximum likelihood, and branches are labelled as foreground for HyPhy analyses. In HyPhy execution, analyses across genes, methods and branch sets are run in parallel, and the random seed, number of processor threads and HyPhy version are recorded for each analysis. In results summary, results are collected into a single table linking each analysis to its gene and branch set. Input files remain unchanged, and decisions made at each stage are recorded in reports.

**Figure 2.**
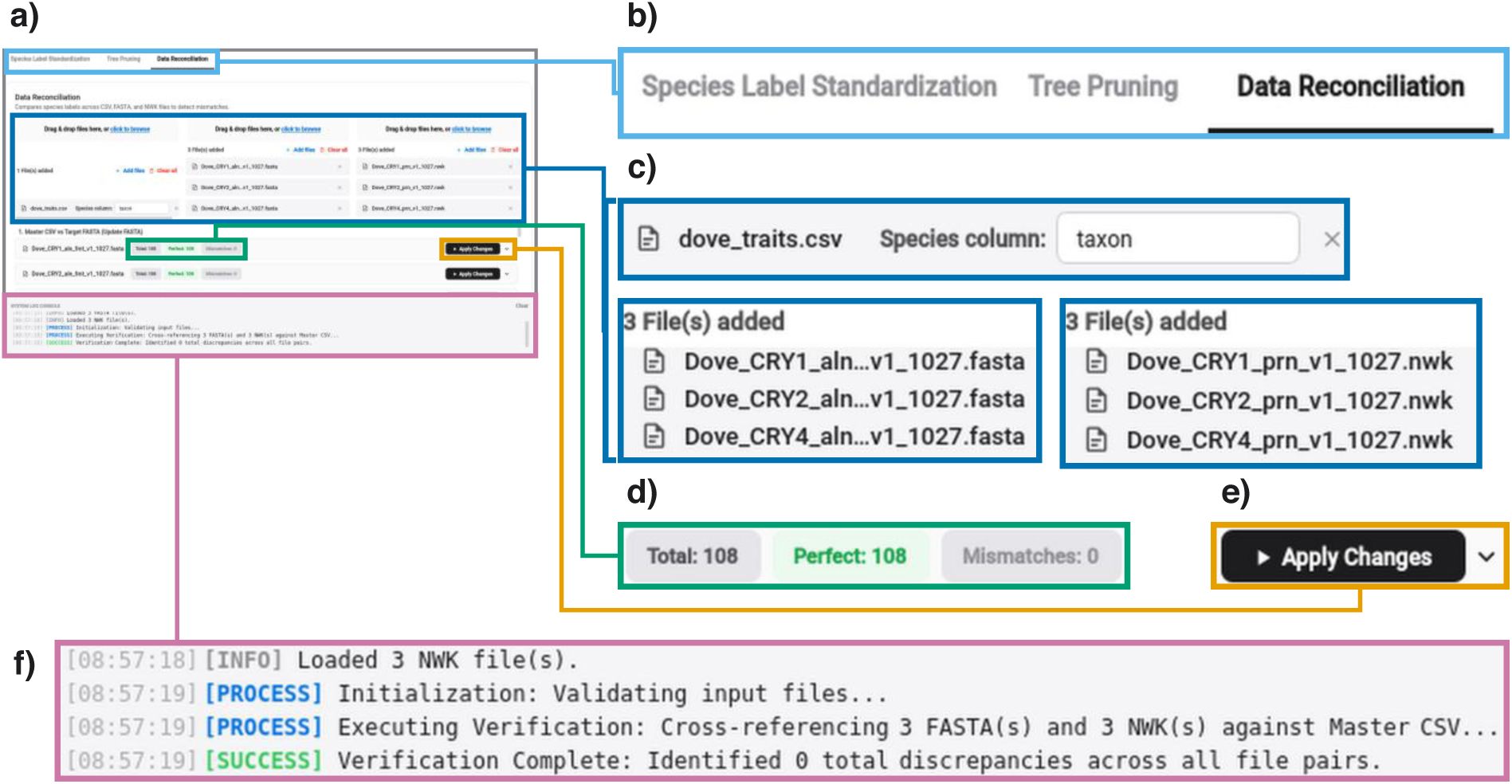
**Data reconciliation in the data preparation stage of HYphlow**, shown with the cryptochrome (CRY) dataset of Sweet et al. (2026) (screenshots of the HYphlow interface). **(a)** The Data Reconciliation subtab. **(b)** The three subtabs of data preparation: species label standardization, tree pruning and data reconciliation. **(c)** Input files: the trait table in comma-separated values (CSV) format with its species column selected, and three codon alignments in FASTA format and three pruned trees in Newick (NWK) format, one for each CRY gene. **(d)** Summary of one comparison between the trait table and an alignment, in which all 108 species labels matched exactly (Perfect: 108; Mismatches: 0). **(e**) Button that applies the selected corrections to the prepared files. **(f)** Log recording each step of the comparison and its outcome. Coloured frames link each enlarged region in **(b)-(f)** to its position in **(a).** Colours within the screenshots are those of the HYphlow interface.

### HYphlow workflow

#### Data preparation

HYphlow takes one or more codon alignments in FASTA format and one or more phylogenetic trees in Newick format, and a trait table in CSV format, all prepared outside the program. HYphlow accepts a shared species tree or gene-specific trees. Species names in the trait table are first compared against a local copy of NCBI Taxonomy using taxopy. Names that exactly match a single NCBI scientific name at the species or subspecies rank are accepted, while close spellings are proposed as corrections. Only scientific names are searched, so synonyms are left for the researcher to review. Names that match several taxa, match only a higher rank, are provisional (e.g., sp., cf.) or are subspecies matched only at the species rank are flagged for review and kept as they are. NCBI Taxonomy serves as a reference for name standardization, while species identity remains the researcher’s decision.

Taxon labels in codon alignments and phylogenetic trees are reduced to species or subspecies names by removing additional information, such as gene symbols and accession numbers. HYphlow compares taxon labels between the alignment and tree of each gene in both directions. When a trait table is provided, its taxon labels are also used as a reference for matching. Identical labels are accepted, while differences are evaluated through prefix matching, followed by string similarity calculated with RapidFuzz. A label found by prefix matching, or the closest label is proposed when its normalized similarity score (RapidFuzz `fuzz.ratio`, 0-100) is at least 80. This fixed threshold was evaluated by simulation. In both the NCBI and cross-file steps, proposed corrections are preselected but remain subject to the researcher’s decision, while unmatched names are retained as they are and the reasons are recorded. HYphlow writes corrected copies of the trait table, alignments and trees, keeping the sequence data and original tree topology. The original and resulting labels, whether each label was accepted automatically, changed to a proposed correction or kept after the researcher declined it, and unresolved mismatches are recorded in Excel reports generated during data preparation.

When a shared species tree is provided, HYphlow uses ETE3 to prune it to the taxa in each alignment, based on identical taxon labels. Pruning preserves phylogenetic distances among the retained taxa. Pruning is recorded in a separate Excel report listing taxa retained in the tree, removed during pruning, or found only in the alignment. A pruned tree is generated when at least three taxa are shared.

#### Tree annotation

Because trait data describe only the taxa at the tips of a tree, HYphlow uses ASR to define foreground branches from trait states. The researcher selects one or more trait columns and the values to designate as the foreground. Taxa carrying any selected foreground value are assigned state 1 (foreground), and the remaining taxa are assigned state 0 (background). HYphlow then uses ASR to assign foreground status to internal branches. Tips with unmatched labels are assigned background and listed in a warning for researcher review. Each branch is assigned the state of its descendant node, so a branch leading to a node reconstructed as foreground is labelled foreground.

HYphlow implements ASR using maximum parsimony (MP) and maximum likelihood (ML) on each tree. ETE3 is used to read the trees and draw the annotated trees. MP uses Fitch parsimony (Fitch 1971) and Hartigan’s method for nodes with more than two descendants (Hartigan 1973). Nodes with multiple equally parsimonious states are recorded as ambiguous. For branch labelling, an ambiguous node takes the state of its parent, while an ambiguous root is assigned background. ML uses a symmetric two-state Markov model with a single transition rate (Mk1) to estimate ancestral state probabilities (Pagel 1999; Lewis 2001). Under ML, a node is assigned foreground when its marginal foreground probability exceeds 0.5. By default, the transition rate is estimated separately for each tree, and a trait whose estimated rate approaches the upper limit of the search range is flagged as having potentially weak phylogenetic signal.

HYphlow generates three annotated trees representing MP, ML, and consensus assignments. Because the two methods can assign different states to the same node, the consensus assigns a branch to the foreground only when both MP and ML reconstruct state 1 and the MP assignment is unambiguous. HYphlow also samples ancestral states 5,000 times under the fitted ML model, conditional on the observed tip states. A branch remains in the consensus foreground only when more than half of the samples assign state 1 to its descendant node. These settings were evaluated by simulation. The researcher chooses which of the three annotated trees to use in HyPhy analyses. From the consensus tree, separate trees can also be saved that contain only branches where the trait changes to foreground, only internal foreground branches, or only terminal foreground branches (Fig. 3).

**Figure 3.**
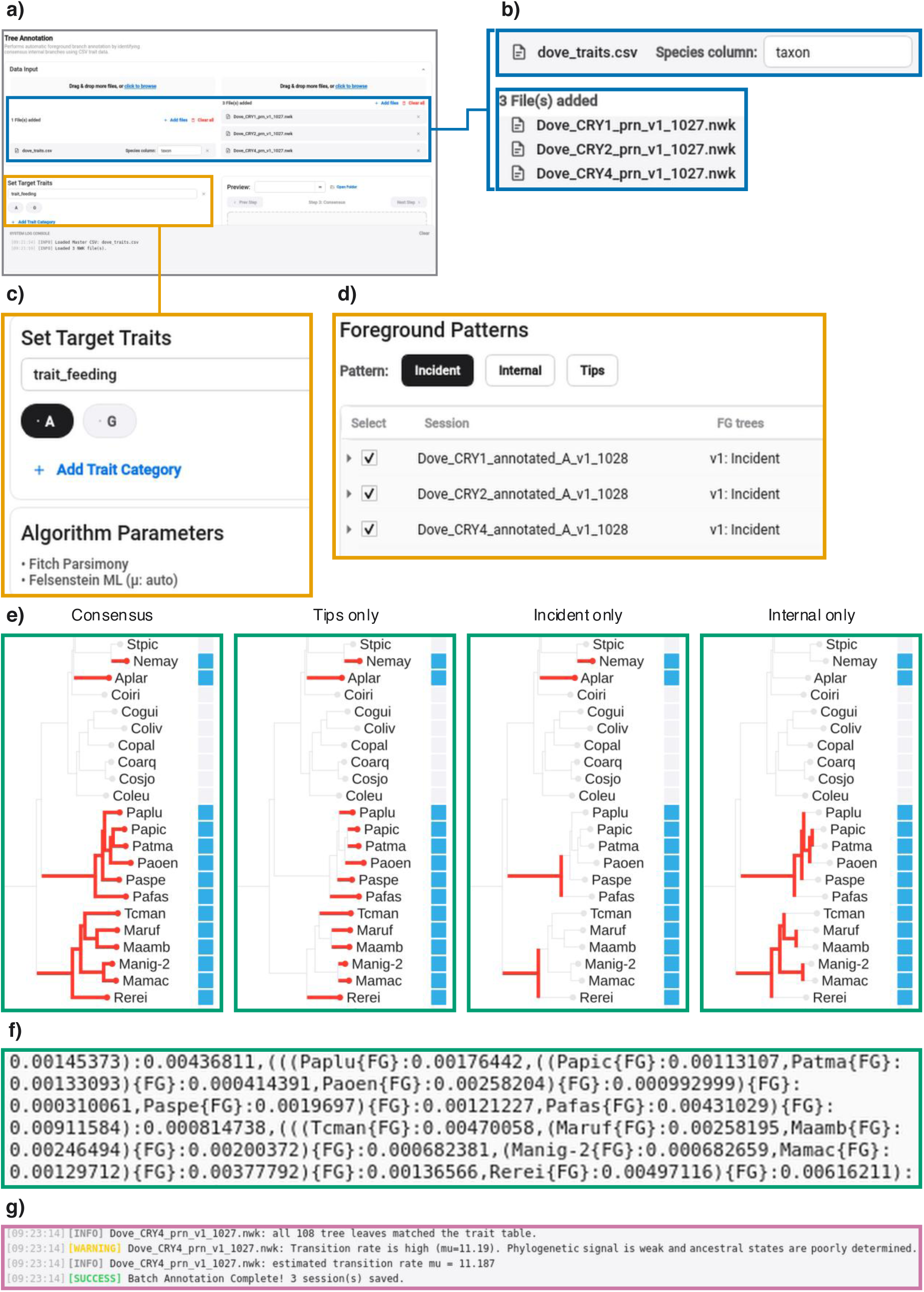
**Foreground branch annotation in the tree annotation stage of HYphlow**, shown with the cryptochrome (CRY) dataset of Sweet et al. (2026) (screenshots of the HYphlow interface). **(a)** The Tree Annotation tab. **(b)** Input files: the trait table in comma-separated values (CSV) format with its species columns selected and three pruned trees in Newick (NWK) format, one for each CRY gene. **(c)** The trait and its foreground state (A, arboreal), and the two ancestral state reconstruction (ASR) methods, Fitch parsimony and Felsenstein maximum likelihood (ML), with the transition rate (*μ*) estimated from the data. **(d)** Foreground pattern selection and the annotated trees saved for each gene. **(e)** The CRY1 tree with consensus assignments and three foreground patterns. The consensus tree shows branches assigned to the foreground by both maximum parsimony and maximum likelihood. The three patterns retain only terminal foreground branches (tips only), branches where the foreground state arises (incident only), or internal foreground branches (internal only). Foreground branches are shown in red, and squares mark taxa in the foreground state. **(f)** Newick text of the consensus tree in (e), with foreground branches labelled {FG}. **(g)** Log reporting the taxa matched between each tree and the trait table, the estimated transition rate and a warning when phylogenetic signal is weak. Coloured frames link enlarged regions to their positions in (a). Colours within the screenshots are those of the HYphlow interface.

Foreground branches are labeled {FG} for HyPhy. An annotation report records the MP and ML states, ambiguity, marginal foreground probability, and foreground assignment for each ancestral node. The transition rate, number of samples, sampling threshold and random seed used for ancestral-state sampling are recorded alongside the report.

#### HyPhy Execution

Selection analyses are run using the command-line interface (CLI) of HyPhy, which can be installed separately or as a dependency of HYphlow. Before each batch, HYphlow locates the HyPhy installation and records its version. HYphlow currently supports six analyses: aBSREL, BUSTED, FEL (Fixed Effects Likelihood), FUBAR (Fast, Unconstrained Bayesian AppRoximation), MEME (Mixed Effects Model of Evolution), and RELAX (Table 2). The analysis options listed in Table 2 are written explicitly into each command, including values that match the HyPhy defaults, so that the recorded settings remain explicit if defaults change between HyPhy versions. HyPhy is run with `--kill-zero-length Yes`, which collapses zero-length internal branches before testing. HYphlow saves the generated commands in a single script that the researcher can review and edit before execution.

**Table 2.** Selection Analyses Run by HYphlow and Their Settings.

| Analysis | Level of inference | Selection tested | Use of foreground branches | HYphlow default settings | Reference |
|---|---|---|---|---|---|
| BUSTED | Gene-wide | Episodic diversifying selection | Tested branches; all branches on unlabelled trees | SRV: Yes; error-sink: No; multiple hits: None | (Murrell et al. 2015) |
| RELAX | Branch set | Relaxation or intensification of selection | Test set, with the remaining branches as reference; runs on labelled trees | SRV:No; models: All; multiple hits: None | (Wertheim et al. 2015) |
| aBSREL | Branch | Episodic diversifying selection | Tested branches; all branches on unlabelled trees | SRV: No; multiple hits: None | (Smith et al. 2015) |
| FEL | Site | Pervasive diversifying or purifying selection | Tested branches; all branches on unlabelled trees | SRV: Yes; multiple hits: None; P-value threshold: 0.1; CI: No; resampling: 0 | (Kosakovsky Pond and Frost 2005) |
| MEME | Site | Episodic diversifying selection | Tested branches; all branches on unlabelled trees | *ω* rate classes: 2; multiple hits: None; impute states: No; P-value threshold: 0.1; resampling: 0 | (Murrell et al. 2012) |
| FUBAR | Site | Pervasive diversifying or purifying selection (Bayesian) | All branches | Grid: 20 x 20; Concentration parameter: 0.5 | (Murrell et al. 2013) |

**Note—**All analyses use the universal genetic code and collapse zero-length internal branches (--kill-zero-length Yes). Every setting listed is written into each HyPhy command, including those equal to the HyPhy defaults. Each run is given a random seed between 10,000 and 99,999 through HyPhy's RANDOM\_SEED parameter (ENV = "SetParameter(RANDOM\_SEED,*n*,0);") and the number of processor threads through CPU = *n*; both are recorded for every run. The *P*-value thresholds for FEL and MEME are those written into the HyPhy commands; significance in the result summary is assessed separately. SRV, synonymous rate variation among sites; error-sink, a model component that accommodates residual alignment errors; multiple hits, instantaneous multinucleotide substitutions; *ω*, ratio of nonsynonymous to synonymous substitution rates; CI, confidence interval; resampling, parametric bootstrap replicates.

Each alignment is paired with its tree, or with alternative annotated trees from different foreground assignments, based on the organism and gene recorded in the project manifest. Before execution, HYphlow compares the taxon labels in each alignment-tree pair and reports mismatched labels, duplicate taxon labels within an alignment and identical sequences. Pairs with mismatched or duplicate taxon labels are deselected by default, and the researcher decides whether to include them. Foreground branches are used as the tested branches in BUSTED, aBSREL, FEL, and MEME, and as the test set in RELAX, with the remaining branches serving as the reference set. On unlabelled trees, BUSTED, aBSREL, FEL and MEME test all branches, whereas RELAX runs only on labelled trees because it requires separate test and reference sets. FUBAR analyzes all branches regardless of foreground labels.

Analyses run in parallel, and the researcher sets the number of concurrent analyses and processor threads assigned to each analysis (one by default). The same analyses can also be exported as a SLURM (Simple Linux Utility for Resource Management) array script for computing clusters. By default, HYphlow runs each analysis once, with a random seed passed to HyPhy and recorded. Alternatively, an analysis is run three times with different seeds, and the three model fits are kept together for comparison. An analysis that stops with an error or writes an empty result is retried once with `TOLERATE_NUMERICAL_ERRORS=1`, and the retry is marked in the run log. The seed, number of threads, HyPhy version, start and end times and outcome of every run are recorded in the corresponding batch folder (Fig. 4).

**Figure 4.**
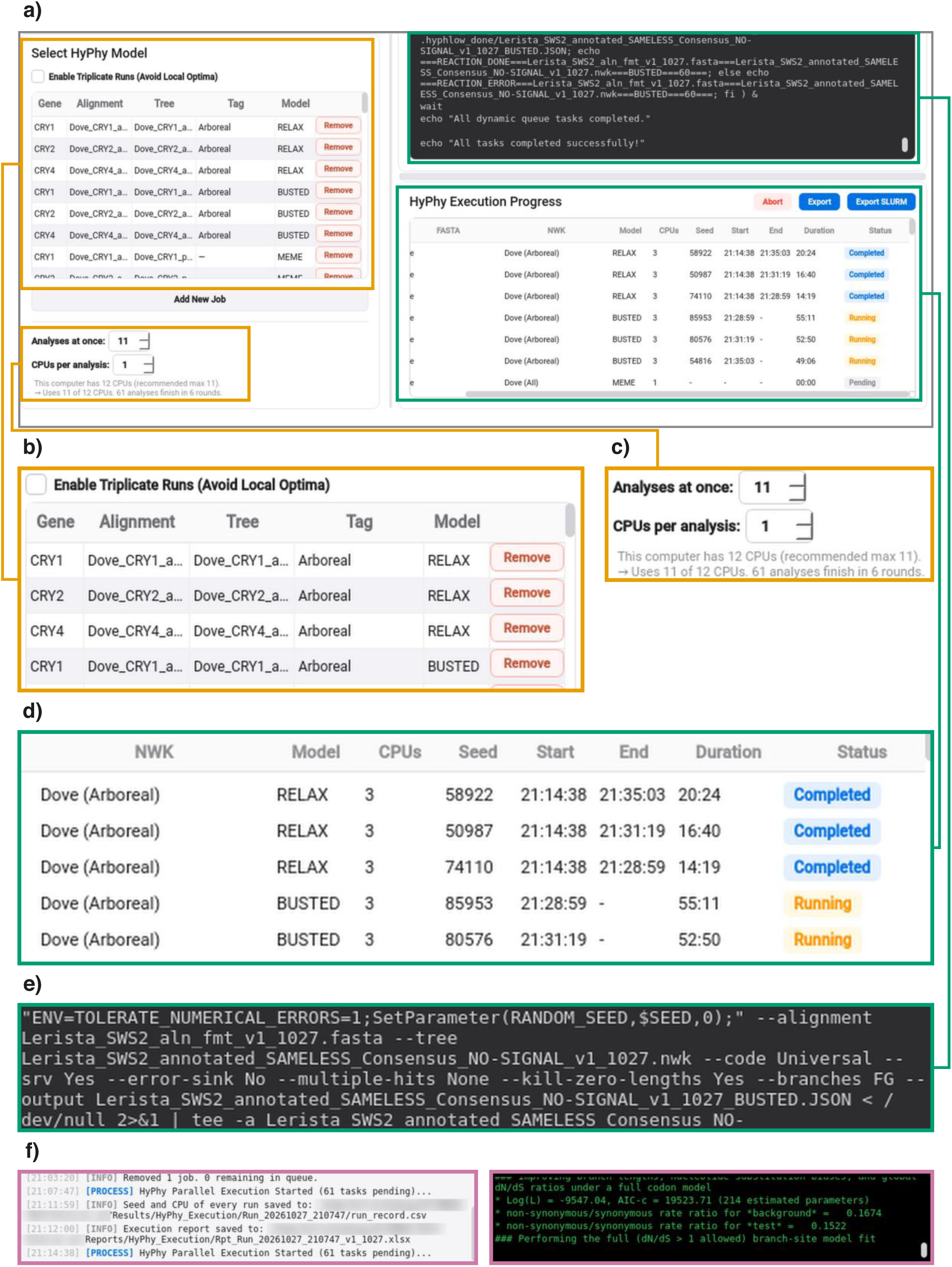
Running HyPhy selection analyses in batches in HyPhy execution stage of HYphlow. (screenshots of the HYphlow interface). **(a)** The HyPhy Execution tab. **(b)** Analyses queued for the run. Each row pairs the alignment of one gene with its annotated tree and the foreground state set in the tree annotation stage. **(c)** The number of analyses run at the same time and the number of central processing units (CPUs) given to each, with the CPUs available on the computer and the number of rounds needed for the queued analyses. **(d)** Progress of each analysis, with its tree and foreground label, selection test, CPUs, the random seed passed to HyPhy, start and end times, duration and status. **(e)** The HyPhy command written for one analysis, specifying the alignment, the tree, the universal genetic code, the foreground branches (`--branches FG`) and the result file. **(f)** Log recording the start of each batch and the files in which seeds, CPUs and the run report were saved (left), beside the HyPhy output for the running analysis (right). Directory names were blurred except for the folders created by HYphlow. Coloured frames link enlarged regions to their positions in (a). Colours within the screenshots are those of the HYphlow interface.

**Figure 5.**
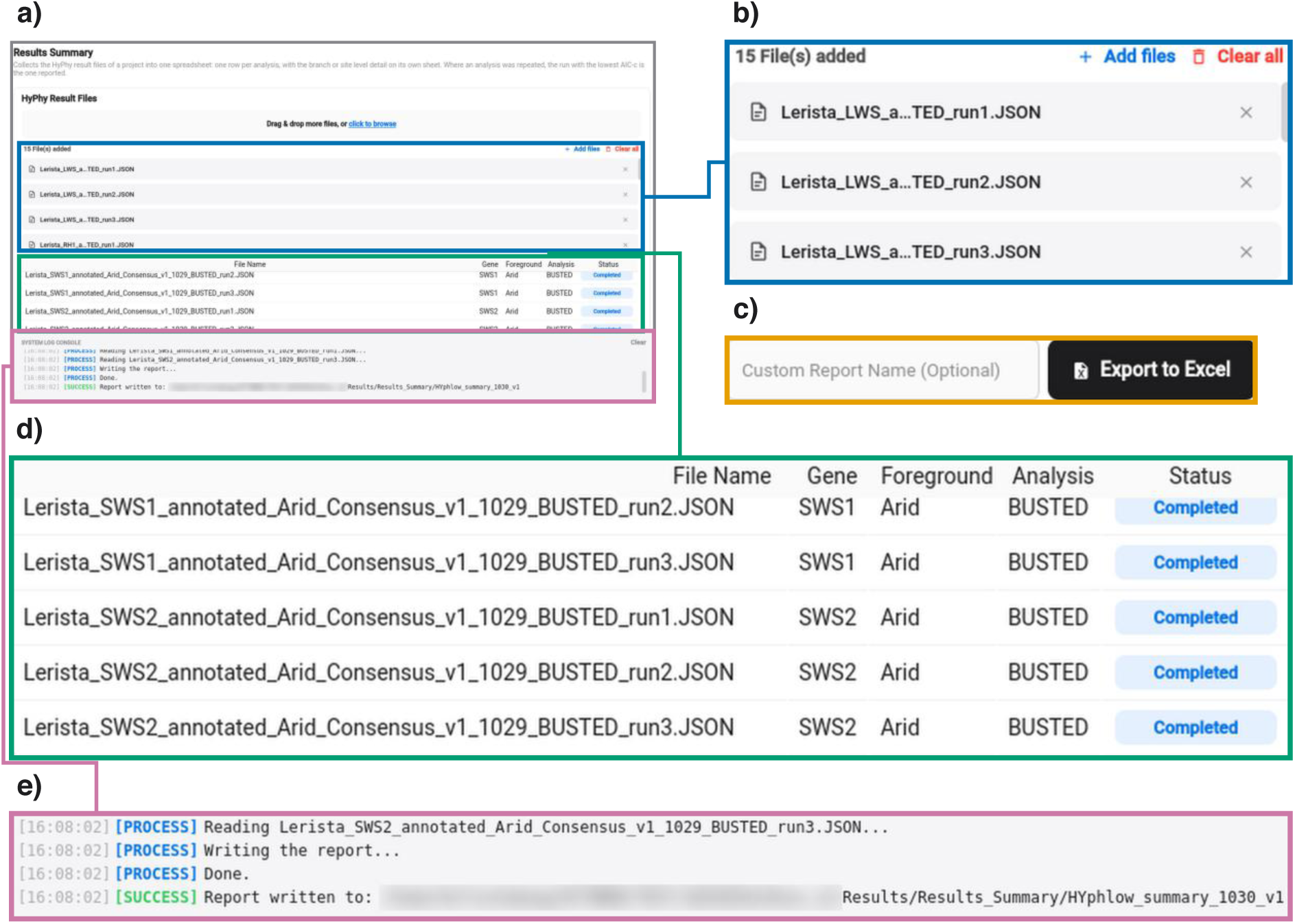
Combining HyPhy result files into one report in the result summary stage of HYphlow. (screenshots of the HYphlow interface). **(a)** The Results Summary tab. **(b)** HyPhy result files in JavaScript Object Notation (JSON) format added to the project: BUSTED results for the five opsins with arid habitat as the foreground state, each analysis run three times (three of 15 files shown). **(c)** The optional report name and the button that saves the report as an Excel workbook, from a separate capture of the row below the log in **(a)**. **(d)** The loaded files, with the gene, foreground state and selection test that HYphlow assigns to each file from its name, and whether each file was read. The workbook contains one row per analysis and separate sheets for branch- and site-level results. When an analysis was run more than once, the run with the lowest small-sample corrected Akaike information criterion (AICc) is reported. **(e)** Log reporting the result files read and the report saved, with the folder in which it was saved. Directory names were blurred except for the folders created by HYphlow. Coloured frames link enlarged regions to their positions in **(a)**. Colours within the screenshots are those of the HYphlow interface.

#### Results summary

Once the HyPhy analyses are complete, HYphlow combines the JSON result files into one spreadsheet, taking all values directly from these files. Replicate runs of the same alignment, tree and analysis are grouped. For BUSTED, RELAX and aBSREL, the run with the lowest small-sample corrected Akaike information criterion (AICc) under the alternative model is reported, and for FEL, MEME and FUBAR, which analyze selection and individual sites, the first run is reported. The AICc of every run is retained, because repeated runs can reach different likelihood optima. For BUSTED and RELAX, HYphlow adds a note when replicate runs differ in significance, in the direction of K or in the number of *ω* rate classes, or when HyPhy records a negative likelihood ratio or a convergence warning.

The summary sheet contains one row per analysis, recording the organism, gene and foreground set. Analyses without foreground labels are recorded as testing all branches. For BUSTED and RELAX, each row gives the P-value of the reported run, along with the highest *ω* class and its proportion of the tested branches. For RELAX, the selection intensity parameter K is also reported, with K > 1 indicating intensification and K < 1 indicating relaxation. The model fits for every run of BUSTED and RELAX, including all *ω* classes for each branch set, are recorded in separate sheets. For aBSREL, FEL, MEME and FUBAR, each row gives the number of branches or sites identified as being under positive selection, and the branches or sites themselves are listed in separate sheets. For FEL, the row also gives the number of sites under negative selection.

Significance is determined using the criterion for each HyPhy analysis: P ≤ 0.05 for BUSTED and RELAX, corrected P ≤ 0.05 for aBSREL, P < 0.1 for FEL, P ≤ 0.1 for MEME and posterior probability ≥ 0.9 for FUBAR. Because BUSTED provides a P-value without a significance label, HYphlow applies the P ≤ 0.05 criterion. An aBSREL, FEL, MEME or FUBAR analysis is classified as significant when at least one branch or site meets its corresponding criterion.

### Evaluation by simulation

We evaluated two stages of HYphlow by comparing their outputs with known or independently obtained reference results. Simulated spelling variants of species names were used to evaluate the corrections proposed during data preparation. Traits simulated on phylogenetic trees with known ancestral states were used to evaluate ASR and foreground branch assignment during tree annotation. HYphlow’s ASR was also compared with those produced by established ASR software. These evaluations assess HYphlow’s matching and branch assignment rules against known outcomes, as well as the agreement of its ASR implementations with independent software.

#### Taxon-label matching evaluation

To evaluate corrections proposed for spelling differences among input files during cross-file reconciliation, we simulated spelling errors using species names obtained from NCBI Taxonomy. Names were sampled randomly from Eukaryota, Fungi, Bacteria, and Viruses, with Fungi also included in Eukaryota sampling pool, and only binomial formats were retained to focus on spelling differences. Reference sets were assembled randomly from each group and contained names from multiple genera, representing the standardized names against which labels are reconciled. Each simulated dataset contained 50 reference names and 10 names excluded from the reference set to represent absent taxa. Using custom Python scripts with fixed random seeds, we generated 10 spelling variants for each reference name at each of six edit levels, applying one to six random single-character insertions, deletions, or substitutions. This produced 3,000 spelling-error comparisons and 10 absent-taxon comparisons per dataset.

Each simulated label was compared against the reference set using HYphlow’s cross-file matching rules. Proposed corrections matching the original species name were classified as correct, and those matching a different reference name as incorrect. Comparisons with no qualifying match were recorded as unresolved. We measured the rates of correct and incorrect corrections for spelling variants and incorrect corrections for absent taxa across similarity thresholds from 50 to 100 in increments of one. Rates were calculated as percentages of all comparisons within each condition, including unresolved comparisons. Because proposed corrections are preselected in HYphlow, the incorrect correction rate measures how often an incorrect match would be applied if the researcher accepted the preselected correction.

For each taxonomic group, the lowest threshold producing no more than 1% incorrect suggestions in both conditions was selected using sequential sampling, with up to 2,000 datasets evaluated per group. Correction rates were calculated from cumulative comparison counts, and sampling stopped when the selected threshold and suggestion rates met predefined stability criteria between consecutive checkpoints or the maximum number of datasets was reached. The threshold selected from Eukaryota was then tested using 1,000 independently generated datasets for each of the four groups, with correction rates recalculated at every threshold. We also evaluated HYphlow’s fixed threshold of 80 on the same independent evaluation datasets. We calculated 95% Confidence intervals using 2,000 dataset-level bootstrap replicates, because comparisons within a dataset share the same reference set. Detailed simulation procedures and additional analyses are provided in Supplementary Note S1.

#### Tree annotation evaluation

Because ancestral states are generally unknown in empirical datasets, the accuracy of these assignments cannot be assessed directly. We therefore simulated phylogenies and binary traits with known histories to evaluate HYphlow’s tree annotation (Table 3). We assessed the agreement of HYphlow’s ASR with established implementations, the accuracy of ASR and foreground branch assignments, and the effects of consensus settings on foreground assignments.

**Table 3.** Simulation design for tree annotation evaluation.

| Component | Simulation settings | Number of datasets |
|---|---|---|
| Tree topology | Balanced, caterpillar, Yule | NA |
| Tree size | 8, 16, 32, 64, 128, and 256 tips | NA |
| Branch lengths | Equal, or drawn from an exponential distribution (rate 1); minimum 10<sup>-4</sup> | NA |
| Tree replicates | 10 per combination (360 trees) | NA |
| Symmetric transitions (SYM) | Equal gain and loss rates (q<sub>01</sub> = q<sub>10</sub> = *μ*); μ = 0.1, 0.5, 2.5, 10 | 1,440 |
| Asymmetric transitions (ARD) | Gain rate four times the loss rate (q<sub>01</sub> = 1.6*μ*, q<sub>10</sub> = 0.4*μ*); same four *μ* values | 1,440 |
| Branch-specific rate variation (RV) | SYM with branch-specific rate multipliers (gamma, shape 1, mean 1); same four *μ* values | 1,440 |
| Single gain (SG) | One gain without subsequent losses | 360 |
| No phylogenetic signal (NS) | Independent tip states; foreground probability of 0.25 or 0.5 | 720 |
| Total simulated datasets | 360 trees × 15 trait conditions | 5,400 |
| Datasets with variable tip states | Used for comparisons with phangorn and castor | 4,810 |
| Known-state evaluation datasets | Variable tip states, excluding NS | 4,097 |

**Note—** Binary traits were simulated on 360 trees under 15 trait conditions. Branch lengths were rescaled to mean root-to-tip distance of 1. Under SYM *μ* represents the expected number of transitions along an average root-to-tip path. NS datasets had no simulated ancestral states and were excluded from accuracy evaluations. Datasets with uniform tip states were excluded from reconstruction analyses. *Abbreviations:* SYM, symmetric transitions; ARD, asymmetric transitions; RV, branch-specific rate variation; SG, single gain; NS, no phylogenetic signal; *μ*, simulated transition rate; NA, not applicable.

We generated trees with different shapes, sizes and branch lengths to evaluate HYphlow under a range of conditions. We also simulated traits with different rates and patterns of evolution. Trees with 8 to 256 tips were generated with three topologies: balanced, caterpillar, and Yule. Balanced and caterpillar trees represented highly balanced and highly unbalanced topologies, respectively, while Yule topologies were generated by a random branching process (Harding 1971; Fischer et al. 2023). Branch lengths were either equal or drawn from an exponential distribution. In total, we generated 360 trees, with 10 replicates for each combination of tree size, topology and branch-length distribution (Table 3). For balanced and caterpillar trees with equal branch lengths, the topology and branch lengths were identical across replicates, and only the species assigned to tips differed. All trees were rescaled to a mean root-to-tip distance of 1, allowing (*μ*) to be compared across trees.

Every dataset could be regenerated using its recorded random seeds, and all 5,400 datasets (360 trees × 15 trait conditions) passed checks covering tree structure, recorded transition histories and the rules of each simulation scenario. None of the pooled statistical diagnostics exceeded the predefined thresholds (Supplementary Note S2.1). Datasets in which all tips had the same state were excluded because they provided no variation in tip states for evaluating ASR. Of the 5,400 datasets, 4,810 were used to compare HYphlow’s reconstructions with those of established R packages. After excluding NS datasets, 4,097 datasets with known ancestral states remained for accuracy evaluation.

We compared HYphlow’s reconstructions with those of established R packages to check their agreement before evaluating reconstruction accuracy. MP reconstructions were compared with those of phangorn v2.12.1 (Schliep 2011), and ML reconstructions with those of castor v1.8.7 (Louca and Doebeli 2018), on the same trees and tip states, using the symmetric two-state model with a flat root prior for ML. Only the root and internal nodes were compared. For MP, we compared the most-parsimonious reconstruction (MPR) state sets, which contain both states at ambiguous nodes. For ML, we compared marginal foreground probabilities under two rate conditions. Each implementation first estimated its own transition rate, as in a standard analysis. For SYM datasets, both then used the simulated rate, separating differences in likelihood calculation from differences in rate estimation. ML probabilities were considered to agree when they differed by no more than 10^-4^. For SYM datasets in which probabilities at the simulated rate differed by more than 10^-4^, castor’s probabilities were recalculated with analytical transition probabilities for the two-state model instead of its default approximation. We also compared the states assigned from these probabilities, with a node assigned to the foreground when its probability exceeded 0.5. Nodes at which either program’s probability was within 10^-4^ of 0.5 were excluded from the state agreement comparison and reported separately as ties, because small numerical differences can change the state assigned to them.

HYphlow searched for transition rates between 0.01 and 100 and reported estimates above 95 as a warning of weak phylogenetic signal. When HYphlow estimated a rate above 95 but castor estimated a lower rate, we calculated the log-likelihood at both estimates using HYphlow’s likelihood function. A log-likelihood more than 10^-6^ higher at castor’s estimate was taken to indicate that HYphlow’s rate search had missed a better-fitting rate.

Reconstructed states were compared with the simulated ancestral states using the original simulated trees, so the accuracy was measured without errors from tree estimation. We evaluated four reconstruction methods: HYphlow MP, phangorn MP, HYphlow ML and castor ML. Only the root and internal nodes were evaluated. MP accuracy was calculated using only unambiguous nodes, with the proportion of ambiguous nodes reported separately. For comparison with ML across the same node ML accuracy was calculated using the state with the higher marginal probability at each node, with each program using its own estimated transition rate. We evaluated ML probabilities using the Brier score (Brier 1950), which accounts for both the predicted probabilities and the simulated ancestral states. Accuracy was reported for each simulation condition, tree size, topology and branch-length distribution, as well as overall. Overall accuracy was calculated from the total number of evaluated nodes across datasets, giving larger trees more weight.

HYphlow’s MP and ML methods assign a branch to the foreground when its descendant node is reconstructed in state 1, with ambiguous MP nodes resolved by a fixed tie-breaking rule that follows the parent’s state when possible and otherwise assigns state 0. The default consensus method assigns a branch to a foreground only when MP assigns an unambiguous state of 1, ML assigns state 1, and more than half of 5,000 ancestral state samples under the fitted ML model support state 1. This method limits foreground assignment to branches supported by both MP and ML. We evaluated HYphlow’s MP, ML and consensus assignments of the 4,097 datasets used for accuracy evaluation. Only internal branches were evaluated, because the root has no incoming branch and tip branches carry the observed state under all three methods. Each internal branch was given the simulated state of its descendant node, following the same rule used by HYphlow. Assignments were evaluated using precision and recall. Precision measures the proportion of branches assigned to the foreground that were truly foreground, whereas missed foreground branches leave simulated foreground branches in the background set. Specificity was also reported, because precision depends on the proportion of foreground branches.

The consensus method has two settings: the threshold for the proportion of sampled ancestral states assigned to the foreground (default 0.5) and the number of ancestral state samples (default 5,000). We refer to branches with an unambiguous MP state of 1 and an ML state of 1 as consensus eligible. The threshold was varied from 0.3 to 0.9 in increments of 0.1, with MP and ML reconstructions and sampled foreground proportions unchanged, to evaluate its effect on precision and recall. Because HYphlow does not fix the random seed for sampling, repeated analyses of the same data can give different consensus assignments. To measure this variation, sampling was repeated on all 4,097 datasets, using 500, 1,000, 2,000, 5,000 and 10,000 samples with 20 random seeds per dataset at the default threshold. MP and ML reconstructions were kept unchanged across these runs. Disagreement was calculated as the proportion of pairs of seeds that assigned different states to the same branch. Before the analysis, we set a target of no more than 1% disagreement across all internal branches. Disagreement was also calculated for consensus eligible branches alone, because only these branches can receive different consensus assignments across runs. We also counted datasets in which at least one branch received different assignments across seeds.

Because nodes and branches from the same tree are not independent, and trees with the same size and replicate number shared species, with the equal and variable trees of each shape sharing one topology, we calculated 95% confidence intervals using 2,000 cluster bootstrap replicates. Datasets with the same tree size and replicate number were treated as one cluster. For each tree size, 10 clusters were sampled with replacement from the 10 available clusters.

Confidence intervals were calculated using the 2.5^th^ and 97.5^th^ percentiles. The same resampled clusters were used for all methods, allowing paired comparisons between methods. Detailed procedures, settings and validation are provided in Supplementary Note S2.

### Application to published data

To apply HYphlow to empirical data, we reanalyzed two published datasets in which foreground branches had been defined in different ways. Sweet et al. (2026) reconstructed ancestral states with a model-based method in R and labelled as foreground only the branches on which a transition into the foreground state was inferred, whereas Ford et al. (2026) selected foreground branches manually. We took the published alignments, trees and trait data from both studies through HYphlow’s data preparation, tree annotation, HyPhy execution and results summary stages.

We used the cryptochrome 4a gene (CRY4a) from Sweet et al. (2026), which was the only gene tested for both feeding habit and movement in the study. The codon alignment for 108 Columbiformes taxa, the maximum-likelihood species tree and the trait table were obtained from the study’s repository (Boyd 2026). Ortholog assembly and species-tree inference were not repeated. The species tree was rooted using four outgroups, which were then removed, following the R script provided in the same repository. The trait table listed feeding (arboreal or ground) and movement (migratory or regional) states under abbreviated taxon codes. Species names from Supplementary Table S3 of Sweet et al. (2026) were added in a separate column. The alignment, tree and trait table all identified taxa using these codes. We therefore used the codes to reconcile taxon labels across files and used the species names only to check names against NCBI Taxonomy. HYphlow compared taxon labels in both directions between the alignment and tree and compared them with the trait table. The tree was then pruned to the taxa in the alignment.

We also reanalyzed the visual opsin genes long-wavelength-sensitive (LWS), rhodopsin 1 (RH1), rhodopsin-like 2 (RH2), short-wavelength-sensitive 1 (SWS1) and short-wavelength-sensitive 2 (SWS2) (Ford et al. 2026). We focused on *Lerista*, which includes multiple independent ecological transitions and in which the source study performed its branch-level selection tests. Gene alignments containing only *Lerista* and the species tree that the source study used for branch-site tests were obtained from its Figshare repository (Ford et al. 2026). Trait states were taken from Supplementary Table S3 of Ford et al. (2026). We pruned the tree to the taxa in each opsin alignment with HYphlow, which gave five trees with the same 39 tips. For the *Lerista* dataset, foreground branches were assigned for each trait hypothesis in two ways: with HYphlow’s consensus annotation and with the foreground labels of the source study, taken from the labelled trees in its repository (Ford et al. 2026). BUSTED and RELAX were applied to each opsin with both annotations using HYphlow’s default settings.

## Results and Discussion

### Consistent taxon labels across alignments, trees and trait tables determine which taxa enter the analysis

Inconsistent taxon names are common in compiled biodiversity records (Zermoglio et al. 2016). In a selection analysis, a taxon whose label differs among the alignment, tree and trait table is treated as two different taxa. As a result, the taxon may not be matched to its trait state during foreground branch assignment or ancestral state reconstruction and may be excluded from the test. Comparing labels by eye becomes less reliable as the number of taxa and datasets grows, because a single missing or transposed letter is easy to overlook. HYphlow compares every label across the three files with the same matching criteria, suggests possible matches and records whether the researcher accepts or rejects each suggestion. We evaluated the accuracy of these suggestions for spelling variants and for names absent from the reference list using simulations across four taxonomic groups. We then applied the same matching criteria to reconcile the alignments, trees and trait tables of two published selection analyses.

#### HYphlow’s matching criteria consistently recovered spelling variants across taxonomic groups

HYphlow first standardizes taxon names in the trait table using NCBI Taxonomy. It then compares labels in the alignment and tree with the standardized names and suggests a possible match when the similarity score is at least 80. Spelling variants and taxa absent from the reference list can both have high similarity scores against reference names, so the threshold determines how well the two are separated. Because HYphlow matches labels among files of the same dataset, we evaluated the threshold by simulating spelling errors in reference lists of species names sampled from NCBI Taxonomy.

In simulations across four taxonomic groups, incorrect suggestions for spelling variants remained below 1% at every threshold tested. Thus, the incorrect suggestion rate for absent taxa determined the selected threshold for each group (Table 4, Fig. 6a). The selected thresholds differed among groups (67 for Eukaryota, 77 for Fungi, 82 for Bacteria and 88-89 for Viruses; Fig. 6b). In separate test datasets for Viruses, a single spelling edit occasionally led to an incorrect suggestion (0.01% of 500,000 comparisons). In addition, three of 3,000,000 virus spelling variants exactly matched a different reference name and were therefore accepted as exact matches. Among the names in the sampling pool, 23.1% of virus names (3,465 of 14,997) had "virus" as the second word, compared with 3.9% of bacterial names (1,079 of 27,907) with "bacterium" as the second word (Supplementary Table S4). Within each group, these names share the same second word and can differ by only a few characters in the first word (e.g., *Ripaslav virus* and *Ripaslac virus*), so a single spelling edit can turn one reference name into another.

**Figure 6.**
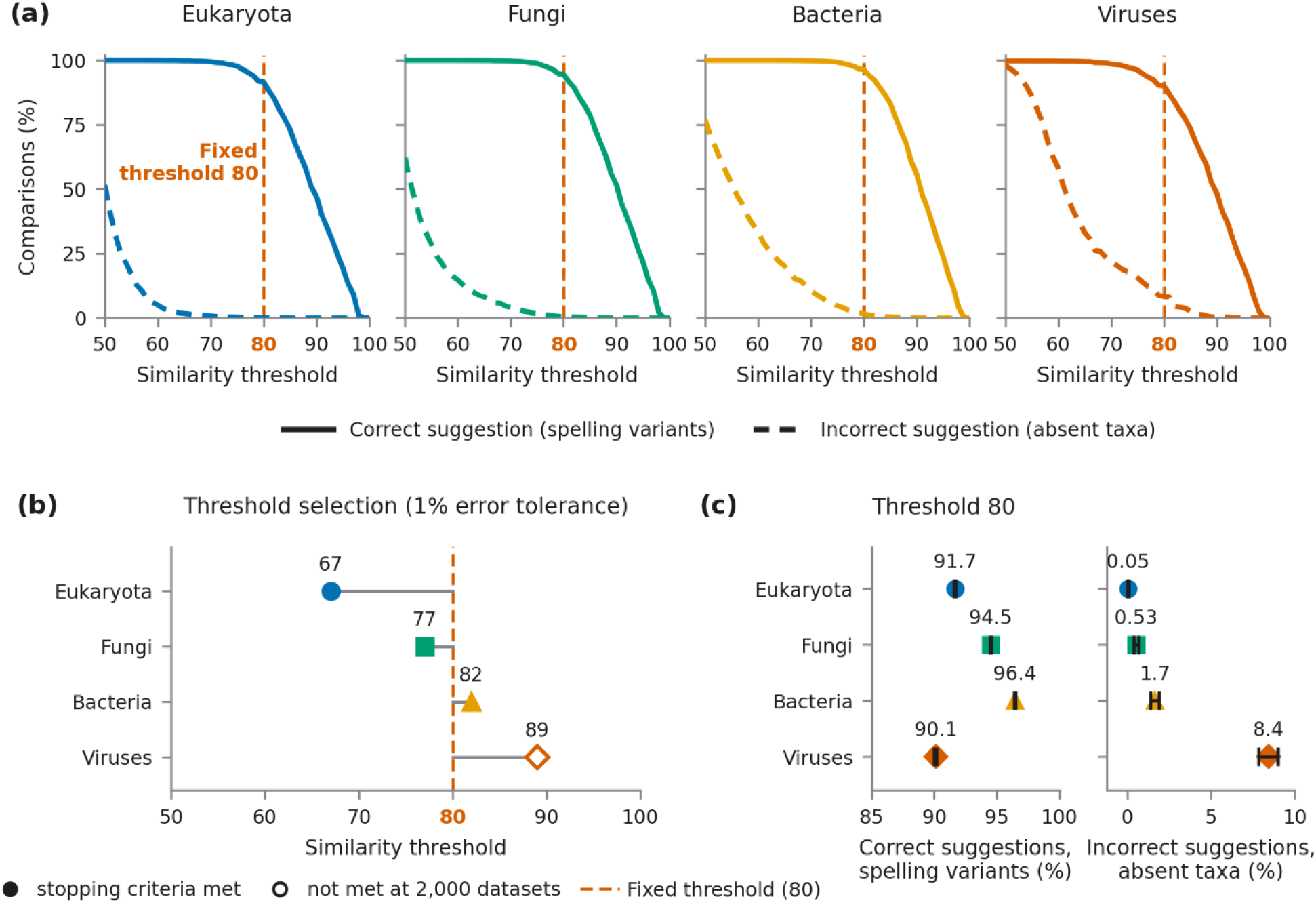
Similarity thresholds that separate spelling variants from absent taxa in four taxonomic groups. **(a)** Percentage of simulated spelling variants that received the correct suggestion (solid lines) and of absent taxa that received an incorrect suggestion (dashed lines) across similarity thresholds from 50 to 100**. (b)** Group-specific thresholds selected by sequential sampling as the lowest thresholds at which incorrect suggestions did not exceed 1% in both spelling variants and absent taxa; grey lines connect each selected threshold to the fixed threshold of 80. Filled symbols indicate that the stopping criteria were met; the open symbol for Viruses indicates that they were not met within 2,000 datasets, and the value shown is from the final checkpoint. **(c)** Correct suggestions for spelling variants and incorrect suggestions for absent taxa at HYphlow’s fixed threshold of 80, with values shown above the symbols. Horizontal bars show 95% confidence intervals from 2,000 data-level bootstrap replicates; most are narrower than the symbols. Groups are distinguished by colour and symbol shape (Eukaryota, blue circles, Fungi, green squares; Bacteria, orange triangles; Viruses, vermilion diamonds). Panels (a) and (c) were calculated from the independent evaluation datasets (1,000 datasets per group; 3,000,000 spelling-variant and 10,000 absent-taxon comparisons per group), and panel (b) from the threshold-selection datasets (Table 4). Details are given in Supplementary Note S1.

**Table 4.** Similarity thresholds selected in four taxonomic groups.

| **Group** | **Eligible names** | **Datasets** | **Threshold** | **Spelling variants** |  | **Absent taxa** |  |
|---|---|---|---|---|---|---|---|
|  |  |  |  | **Corr. (%)** | **Incorr. (%)** | **Corr. (%)** | **Incorr. (%)** |
| Eukaryota | 769,353 | 1,500 | 67 | 99.7 | 0.0001 | 99.02 | 0.98 |
| Fungi | 68,242 | 1,500 | 77 | 97.3 | 0.0004 | 99.19 | 0.81 |
| Bacteria | 27,907 | 1,500 | 82 | 92.4 | 0.0007 | 99.15 | 0.85 |
| Viruses | 14,997 | 2,000 | 89<sup>a</sup> | 52.6 | 0.0076 | 99.45 | 0.55 |

**Note—**Values are from the threshold-selection datasets at the final checkpoint of each group; values from the independent evaluation datasets are given in Supplementary Table S3. Each dataset contained 3,000 spelling-variant comparisons and 10 absent-taxon comparisons, and rates are percentages of all comparisons in each condition. Eligible names were species-rank scientific names in two-word names from NCBI Taxonomy (Supplementary Table S1); Fungi is a subset of Eukaryota. Thresholds were selected as the lowest value at which incorrect suggestions were 1% or fewer in both conditions. For spelling variants, a correct suggestion was the variant's own reference name, and an incorrect suggestion was a different reference name; variants counted in neither column received no suggestion at or above the threshold or exactly matched a different reference name. For absent taxa, any suggestion was incorrect, and correct means that no suggestion was made. Corr., correct; Incorr., incorrect. <sup>a</sup>Provisional: the selected threshold alternated between 88 and 89, and the stopping criteria were not met within 2,000 datasets.

When the Eukaryota threshold (67) was applied to all four groups, it gave a correct suggestion for 99.2-99.9% of spelling variants in the separate test datasets. However, incorrect suggestions for absent taxa increased from 1.10% in Eukaryota to 5.77% in Fungi, 15.01% in Bacteria and 26.15% in Viruses (Supplementary Table S3; Fig. 6a). HYphlow’s fixed threshold of 80 falls within the range of group-specific thresholds (67–89).

At the fixed threshold of 80, 90.1-96.4% of spelling variants received a correct suggestion across the four groups, and fewer than 0.1% received an incorrect suggestion (Supplementary Table S3; Fig. 6c). Spelling variants that received no suggestion (3.6-9.8%) were flagged for review. Among variants with one or two edits, at least 99.95% received a correct suggestion in every group. Incorrect suggestions for absent taxa were 0.05% in Eukaryota and 0.53% in Fungi, but higher in Bacteria (1.66%) and Viruses (8.42%).

#### Label reconciliation in two published selection analyses

In the *Lerista* opsin dataset of Ford et al. (2026), 37 of the 39 species names in the trait table matched NCBI Taxonomy exactly, and none required a spelling correction. The other two names, *Lerista microtis* "east" and "west", included population labels. Both were matched to the species *L. microtis* in NCBI Taxonomy with a similarity score of 82.1, which was lowered only by the population labels, and were flagged for review. We kept the two population labels unchanged to compare them with the labels in the alignments and trees. When we reconciled the labels across files, the five opsin alignments and their pruned trees shared the same 39 labels. Both the alignments and the trees showed the same result against the trait table in all five genes: 36 of 39 labels matched exactly, and three differed (*L. arenicola, L. cinerea*, and *L. microtis*).

To determine why the three sets of labels differed, we examined the supplementary tables of the source study. *L. microtis* "east" and "west" appeared only in the trait table, as did *L. hobsoni*, whereas *L. arenicola* and *L. cinerea* appeared only in the tree and alignment files. Using the museum specimen numbers reported in Supplementary Table S2 of Ford et al. (2026), we found that *L. microtis* "east" was originally catalogued as *L. arenicola* in the South Australian Museum (SAMA; R64970) and that *L. microtis* "west" was catalogued as *L. microtis* (R29496). *L. arenicola* was formerly treated as a separate species and is now included in *L. microtis* (Farquhar et al. 2024), and *L. hobsoni* was formerly included in *L. cinerea* (Couper et al. 2016). The label differences therefore reflected taxonomic revisions rather than spelling differences. We did not accept the suggested matches and edited only the trait table to match the tree and alignment labels. When we repeated the reconciliation, all 39 labels matched exactly across all five genes, and all 39 taxa were included in the analysis with matching trait states.

In the Columbiformes CRY4a dataset of Sweet et al. (2026), the alignment, tree and trait table identified taxa using the same abbreviated codes, and all 108 labels matched exactly across the three files. Because the codes were used as taxon labels, species names were checked against NCBI Taxonomy separately from the labels used in the analysis. Of the 108 species names, 99 matched NCBI Taxonomy exactly. Four names differed from the names in NCBI Taxonomy in spelling or Latin endings (*Hemiphaga novaseelandiae*, *Phapitreron amethystina, Stretopelia decaocto* and *Treron seiboldi*) and were flagged with similarity scores ranging from 90.3 to 98.0. The remaining five names were not found in NCBI Taxonomy and were kept unchanged. All four flagged names had similarity scores above the fixed threshold of 80 used for label reconciliation. These differences in species names did not affect the reconciled alignment, tree or trait table.

#### HYphlow records label differences across files for the researcher to resolve before selection tests

Algorithms developed for scientific names, such as Taxamatch, compare genus names and specific epithets separately and suggest candidate matches from taxonomic reference lists (Rees 2014; Grenié et al. 2023). Taxamatch is designed to find matching scientific names in taxonomic reference lists, whereas HYphlow uses NCBI Taxonomy to standardize names before comparing labels across input files. Comprehensive Automated Pipeline using HyPhy for Evolutionary Inference with Nextflow (CAPHEINE), another workflow for HyPhy analyses, starts with unaligned sequences. It infers gene trees from sequence alignments and identifies foreground branches using a list of pattern of sequence identifiers (Verdonk et al. 2026). Because the trees are inferred from the alignments, the same sequence labels are used in both files, and no separate trait table is required for foreground assignment. HYphlow instead accepts separately prepared alignments, trees and trait tables and reconciles taxon labels across these files.

In both case studies, names with similarity scores above 80 were flagged with suggested matches. In the Columbiformes dataset, the flagged names differed from names in NCBI Taxonomy only in spelling or Latin endings, consistent with the simulations. In the *Lerista* dataset, suggested matches above the same threshold also arose from population labels and taxonomic revisions, and recording which file contained each label showed where corrections were needed. HYphlow therefore provides suggested corrections for researchers to confirm and records each decision in the reconciliation report, allowing corrections made before analysis to be traced. Without reconciliation, three tips of the *Lerist*a tree would not have been matched to a trait state. Because taxon names can differ across data sources for reasons other than spelling, checking labels across files before foreground branch assignment or ancestral state reconstruction prevents taxa from being left without trait state and excluded from the test.

### Foreground branch assignment determines which lineages a selection test treats as the test set

Ancestral state reconstruction can differ between methods, and MP and ML can assign different states to the same node (Cunningham et al. 1998; Joy et al. 2016). In a selection analysis, the reconstructed states determine which branches are labelled as foreground. As a result, the same tree and trait data can lead a selection analysis to compare different sets of lineages, depending on the reconstruction method and on which branches are labelled. Studies also differ in this choice, labelling only branches on which a transition into the foreground state is inferred or every branch in the foreground state, and branches labelled by hand or in a separate program are not always accompanied by the reconstructed states. HYphlow labels a branch as foreground only when MP and ML reconstructions agree on the foreground state, records the reconstructed states and branch assignments, and lets researchers keep either all consensus foreground branches or subsets such as the branches on which the foreground state begins. We compared HYphlow’s reconstructions with those of established implementations using simulated datasets.

#### HYphlow’s reconstructions match established implementations

Reconstruction methods can also disagree, because parsimony minimizes the number of changes, whereas likelihood accounts for branch lengths and estimates transition rates. HYphlow reconstructs ancestral states by MP and ML on each prepared tree. The consensus retains branches for which both methods assign the descendant node to the foreground, the MP state is unambiguous, and more than half of the sampled ancestral-state histories support the foreground assignment. Researchers can retain all consensus foreground branches or subsets containing only incident, internal, or terminal foreground branches. Branches that do not meet these conditions remain in the background and are recorded in the report. How well the consensus rule identifies foreground branches depends on accuracy of each reconstruction, which we assessed by simulation.

HYphlow and phangorn returned identical MPR state sets, including ambiguous states, at all 417,886 nodes across the 4,810 datasets (Supplementary Table S7). When the simulated transition rate was used for SYM datasets, HYphlow and castor assigned the same state at 108,700 of 108,701 nodes, excluding 2,536 ties. The only disagreement occurred at a node where HYphlow assigned a foreground probability of 0.5007, and castor assigned 0.4993 (Supplementary Table S8). At the simulated rate, marginal probabilities differed by no more than 10^-4^ at 96.4% of nodes. For the 541 datasets with larger differences, recalculating castor’s probabilities using analytical transition probabilities reduced the maximum differences to 6.1 × 10^-6^, below the 10^-4^ agreement threshold. This indicates that the differences above the threshold, including the single disagreement in assigned state, arose from castor’s default approximation of branch transition probabilities.

When each program estimated its own transition rate, HYphlow and castor assigned the same state at 385,783 of 385,847 nodes (99.98%), excluding 32,039 ties (7.7% of all nodes). Ties were more common in datasets without phylogenetic signal and in datasets with high transition rates (Supplementary Table S8). Marginal foreground probabilities differed by no more than 10^-4^ at 91.3% of nodes. For datasets in which HYphlow estimated a transition rate of 95 or lower, the two rate estimates differed by less than 1% of castor’s estimate in 85.5% of datasets.

HYphlow estimated rates above 95 and issued a weak phylogenetic signal warning in 510 of the 4,810 datasets (10.6%), including 224 of the 713 NS datasets (31.4%). Castor’s estimates for these datasets ranged from 2.9 to 17,096. In 23 datasets, HYphlow estimated a rate above 95 while castor estimated a rate below 95. The log-likelihoods at the two estimates differed by less than 10^-6^ in 22 of these datasets, so the data did not distinguish between the two rates. In the remaining dataset, the log-likelihood was 0.007 higher at castor’s estimate (2.9), indicating that HYphlow’s rate search had missed a better-fitting rate. How often this warning is triggered is likely to depend on the branch-length scale of the input tree. In our simulations, trees were scaled to a mean root-to-tip distance of 1, so symmetric transition rate of 95 corresponded to 95 expected changes along an average root-to-tip path.

MP reconstructions were ambiguous at 5.8% of the 358,175 root and internal nodes. Among unambiguous nodes, 91.9% (95% CI 91.8-92.1%) matched the simulated ancestral states for both HYphlow and phangorn (Fig. 7a; Supplementary Table S9). Counting ambiguous nodes at half correct gave an MP accuracy of 89.5%. ML accuracy was 90.6% for HYphlow and 90.4% for castor (paired difference 0.22 percentage points, 95% CI 0.18-0.27 percentage points). Their Brier scores were 0.0647 and 0.0648, respectively (Supplementary Table S10). The difference in ML accuracy between HYphlow and castor was no greater than 1.16 percentage points across individual simulation conditions and tree sizes, with the largest difference observed in 8-tip trees.

**Figure 7.**
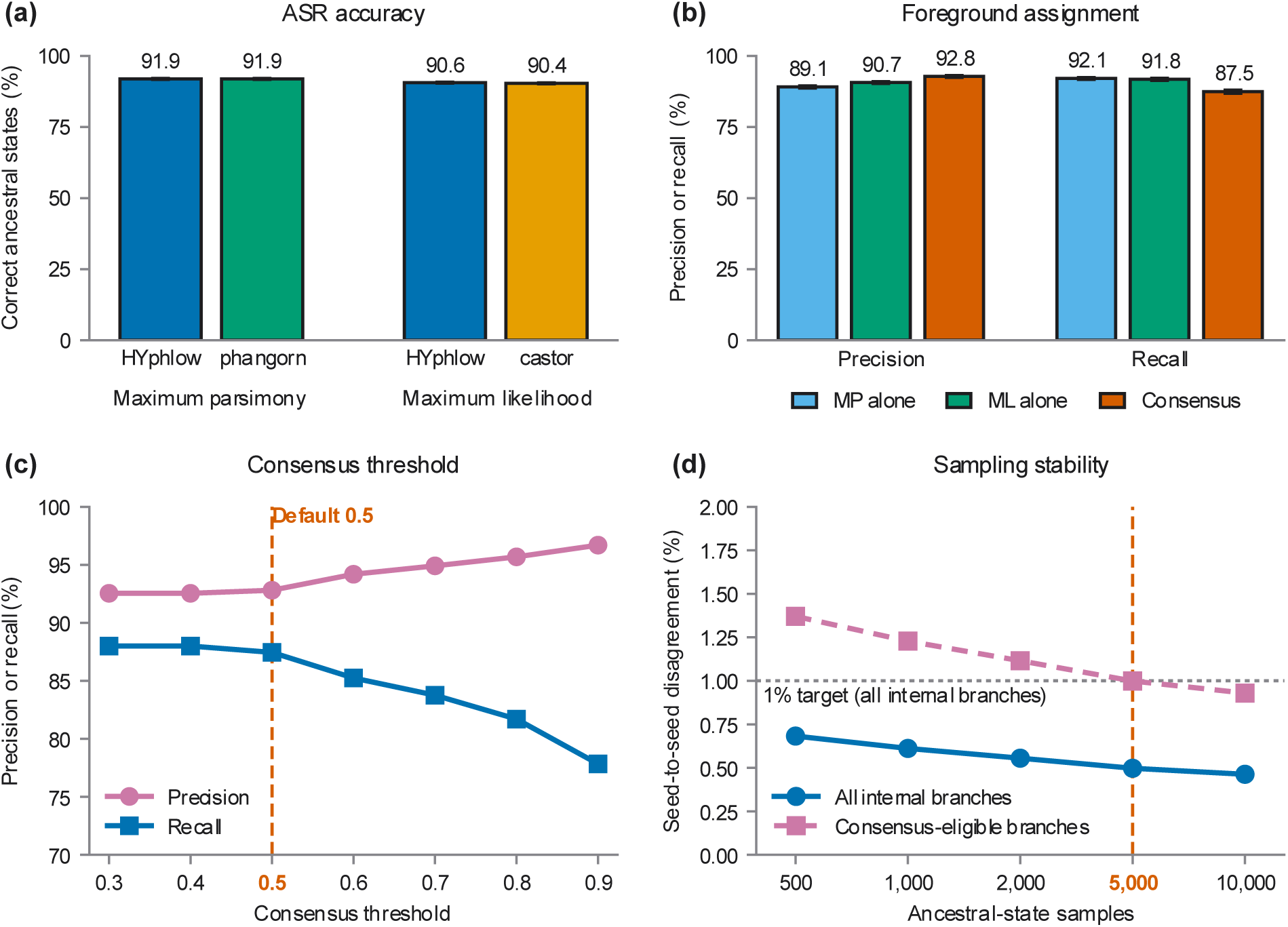
Simulation-based evaluation of HYphlow tree annotation. **(a)** Accuracy of ancestral state reconstruction (ASR) by maximum parsimony (MP) and maximum likelihood (ML) at 358,175 root and internal nodes from 4,097 datasets with simulated ancestral states. HYphlow is compared with phangorn v2.12.1 (Schliep 2011) for MP and with castor v1.8.7 (Louca and Doebeli 2018) for ML. For ML, each program used its own estimated transition rate. HYphlow and phangorn returned identical most-parsimonious state sets at every node. MP accuracy was calculated using nodes with a single most-parsimonious state (94.2% of nodes), whereas ML accuracy was calculated using all nodes. The accuracies shown are therefore directly comparable within each method, but not between MP and ML. **(b)** Precision and recall of foreground branch assignment at 354,078 internal branches using HYphlow’s MP, ML and consensus methods. **(c)** Precision and recall of consensus assignments at consensus thresholds from 0.3 to 0.9 in increments of 0.1. Thresholds of 0.3 and 0.4 produced identical results because no consensus eligible branch had a sampled foreground proportion greater than 0.3 and at most 0.4. The vertical dashed line indicates the default threshold of 0.5. **(d)** Seed-to-seed disagreement in consensus assignments with 500, 1,000, 2,000, 5,000 and 10,000 ancestral state samples and 20 random seeds per dataset. Seed-to-seed disagreement was calculated as the proportion of seed pairs that assigned different states to the same branch. The solid line represents all internal branches and the dashed line represents consensus-eligible branches, defined as branches assigned to the foreground by both an unambiguous MP reconstruction and ML. The horizontal dotted line indicates the predefined 1% disagreement target, which applies to all internal branches. The vertical dashed line indicates the default of 5,000 samples. In **(a)** and **(b)**, error bars denote 95% confidence intervals from 2,000 cluster bootstrap replicates, with clusters defined by tree size and replicate number.

Accuracy decreased as the transition rate increased. At *μ* = 0.1, both methods reconstructed at least 99.8% of nodes correctly in SYM, ARD and RV datasets. At *μ* = 10, MP accuracy, with ambiguous nodes counted as half correct, decreased to 67.2% for SYM, 72.5% for RV and 83.8% for ARD. HYphlow’s ML accuracy decreased to 70.1%, 74.9% and 84.3%, respectively. The higher accuracy in ARD datasets may partly reflect differences in state frequencies. At *μ* = 10, about 80% of tips were in foreground state in ARD datasets, compared with about 50% in SYM datasets. In SG datasets, all unambiguous MP reconstructions matched the simulated states, as expected for a single gain without losses.

Accuracy also increased with tree size, from 77.0% for MP (ambiguous nodes counted as half correct) and 75.3% for ML at 8 tips to 91.1% and 92.1% at 256 tips. However, more datasets with uniform tip states were excluded from smaller trees. As a result, the remaining datasets had different distributions of transition rates across tree sizes. Caterpillar trees had higher ML accuracy than balanced trees (97.4% vs. 86.8%). Caterpillar trees also had shorter total branch lengths and fewer simulated transitions (Supplementary Table S6). Because state frequencies and transition counts differed among scenarios, we could not determine how much of the difference in accuracy was due to using a symmetric model for ARD, RV and SG datasets.

Of the 354,078 internal branches, 52.4% were simulated foreground branches. Consensus had a precision of 0.928 (95% CI 0.925-0.931), compared with 0.891 for MP and 0.907 for ML. However, consensus had a lower recall of 0.875 (95% CI 0.869-0.880), compared with 0.921 for MP and 0.918 for ML (Fig 7b). Consensus also had the highest specificity (0.926), compared with 0.876 for MP and 0.896 for ML. It incorrectly assigned 12,558 simulated background branches to the foreground, compared with 20,847 for MP and 17,536 for ML. HYphlow’s consensus assignment is therefore intentionally conservative: it prioritizes reducing false foreground assignments at the cost of leaving some true foreground branches in the background.

#### Selection analyses across foreground hypotheses are run and summarized together

Applying HyPhy to many genes and foreground hypotheses often requires study-specific scripts to pair alignments with trees, run analyses and collect results. HYphlow brings these steps into one workflow. Once the trees are annotated, HYphlow pairs alignments with trees, runs the selected analyses and collects the results in a single table. The commands, settings and diagnostics are retained for each analysis.

#### HYphlow extends end-to-end HyPhy workflows to comparative studies starting from trait data

Existing tools each cover part of this workflow. The HyPhy command line, Datamonkey and the Galaxy HyPhy tools run individual analyses, HyPhy Vision displays their results, and label-tree and phylotree.js assign foreground branches from taxon lists or by hand. CAPHEINE links these steps for pathogen genomes, carrying unaligned sequences through gene-tree inference, foreground labelling from sequence identifiers and summarized HyPhy results. HYphlow extends this end-to-end approach to comparative studies that begin with an alignment, a tree and trait data, matching species names across the three files, inferring foreground branches from trait states and testing several foreground hypotheses on the same gene.

## Conclusions

HYphlow provides a reproducible framework for testing hypotheses of trait-associated selection across genes and species. It integrates independently prepared sequence alignments, phylogenetic trees, and trait data into a traceable workflow from taxon reconciliation and foreground assignment through HyPhy execution and results summary. Across simulated reference sets from four taxonomic groups, a single similarity threshold effectively distinguished spelling variants from absent taxa, while ancestral-state reconstructions closely matched established implementations. The consensus approach produced more conservative foreground assignments, reducing the assignment of background branches to the foreground. In two published datasets, HYphlow identified inconsistencies among input files before selection analyses, including differences resulting from taxonomic revisions, and accommodated multiple foreground hypotheses for the same gene. Recording taxon reconciliation, ancestral-state reconstruction, branch assignment, and analysis settings allows each selection result to be traced back through the decisions that produced it. Cases in which ancestral states were poorly determined were flagged and recorded alongside the results. Future development could incorporate additional models of trait evolution, including asymmetric and hidden-rate models, and extend evaluation to a wider range of traits and taxonomic groups. Improving the repeatability of HyPhy optimization across replicate runs also remains an important goal. Analyses can be executed locally or exported for computing clusters, allowing the same workflow to scale to larger sets of genes HYphlow is freely available at https://github.com/Schott-Lab/HYphlow/tree/main and can be installed with pip or Bioconda.

## Acknowledgements

Development of this software was supported by an NSERC Discovery Grant to RKS.

