## Supplementary Note for "HYphlow: bringing end-to-end flow to HyPhy selection analyses, from taxon reconciliation to trait-based foreground assignment"

### Supplementary Text

This Supplementary Text provides additional methods and results for the analyses described in the main text. Section S1 describes the simulation used to evaluate how HYphlow matches taxon labels across alignments, trees and trait tables, with the methods followed by the corresponding results. Supplementary tables are numbered consecutively throughout this document.

#### Supplementary Text S1. Evaluation of taxon-label matching on simulated spelling variants and absent taxa

##### *S1.1. Species-name pools*

Species names were obtained from the NCBI Taxonomy database downloaded on October 9, 2026. Scientific names at species rank were collected separately for Eukaryota (taxid 2759), Fungi (4751), Bacteria (taxid 2), and Viruses (taxid 10239). Fungi formed its own pool, and fungal names also remained in the Eukaryota pool. Names were restricted to the binomial form, a capitalized genus name followed by a lower specific epithet (`^[A-Z][a-z]+ [a-z]+$`), so that all simulated labels shared one format. Each name was converted to a taxon label by replacing the space with an underscore, as in HYphlow, and identical labels were kept once within each pool.

The binomial criterion retained 42.6% of species-rank names in Eukaryota, 33.7% in Fungi, 5.0% in Bacteria and 15.6% in Viruses. One duplicated label was removed from Eukaryota (**Supplementary Table S1**).

**Supplementary Table S1**. **Species-name filtering and sampling pool sizes.** Species-rank scientific names were extracted from NCBI Taxonomy and filtered using the binomial-format criteria. Identical name strings were retained only once within each group. The final sampling pools were used to construct simulation datasets.

| Taxonomic group | Species-rank names | Binomial-format names | Duplicated removed | Final sampling pool |
|---|---|---|---|---|
| Eukaryota | 1,806,629 | 769,354 | 1 | 769,353 |
| Fungi | 202,575 | 68,242 | 0 | 68,242 |
| Bacteria | 555,736 | 27,907 | 0 | 27,907 |
| Viruses | 95,848 | 14,997 | 0 | 14,997 |

#### *S1.2. Simulation datasets and label comparisons*

For each simulation dataset, 60 distinct names were randomly selected without replacement from the corresponding pool. 50 names formed the reference set, while the remaining 10 were excluded from this set to represent taxa without a matching reference name. Names could be selected again across different datasets, and all random sampling and spelling-error generation were performed using fixed random seeds for reproducibility.

#### *S1.3 Correction-suggestion evaluation*

Each of the 50 reference names was modified by one to six single-character edits, with 10 variants generated per name and number of edits (**Supplementary Table S*2***). Variants identical to the original name were regenerated, with up to 100 additional attempts. Variants that exactly matched another reference name were retained and recorded as accidental exact matches. Each variant was compared against the 50 reference names, producing up to 3,000 spelling-error comparisons per dataset.

The 10 names excluded from the reference set were compared against the same 50 reference names without spelling modifications. Because their correct names were absent from the reference set, any suggested correction was classified as incorrect, producing 10 absent-taxon comparisons per dataset.

Simulated labels were compared with the reference set using HYphlow's name matching rules. Reference labels were identified through exact matching, followed by prefix matching and RapidFuzz similarity scoring. Suggestions with scores at or above the candidate threshold were classified as correct if they matched the original species name and incorrect if they matched a different reference name. Comparisons without a qualifying suggestion were recorded as unresolved. Accidental exact matches to other reference names were recorded separately.

Correct- and incorrect-suggestion rates were calculated for spelling-error comparisons, whereas only incorrect-suggestion rates were calculated for absent-taxon comparisons:

$$\text{Suggestion rate (\%)} = \frac{\text{Number of suggestions in a given category (n)}}{\text{Total number of comparisons in that condition (N)}} * 100$$

where (n) is the number of correct or incorrect suggestions in the specified category and (N) is the total number of comparisons in that condition, including those without a suggestion.

**Supplementary Table S2. Character-edit operations used to simulate spelling errors.** Each edit type was selected with equal probability, and edit positions were selected randomly, without considering common typing patterns or keyboard proximity. Inserted and substituted characters were drawn from the 26 lowercase letters. Deletion was replaced by insertion when the current label contained four or fewer characters.

| Edit type | Operation | Example |
|---|---|---|
| Substitution | Replace one character with a random lowercase letter | *Panthera_leo* → *Panthera_le**x*** |
| Deletion | Remove one character | *Panthera_leo* → *Panthera_le* |
| Insertion | Insert a random lowercase letter | *Panthera_leo* → *Panthera_le**x**o* |

#### *S1.4 Sequential sampling and threshold selection*

Candidate similarity thresholds ranged from 50 to 100 in increments of one. The lowest threshold with incorrect-suggestion rates of 1% or less in both the spelling error and absent-taxon conditions was selected. This limit was defined before the simulation as a practical tolerance for incorrect suggestions rather than a statistically derived optimum.

To assess the stability of threshold selection, simulation datasets were added cumulatively, and the selected threshold and suggestion rates were recalculated at successive checkpoints. The initial checkpoints were 100, 200, and 300 datasets. Stability was assessed by comparing each checkpoint with the preceding one. The criteria required an unchanged threshold, a difference of less than one percentage point in the correct-suggestion rate, and differences of less than 0.2 percentage points in each incorrect-suggestion rate.

When these criteria were met, subsequent checkpoints were set at 1,000, 1,500, and 2,000 datasets. Otherwise, datasets were added in increments of 100. Sampling stopped when the stability criteria were met across two consecutive comparisons at the larger checkpoints or when 2,000 datasets had been evaluated.

The threshold selected from the Eukaryota datasets was evaluated using 1,000 independently generated datasets for each of the four taxonomic groups. Correction-suggestion rates were evaluated at the selected threshold and at additional thresholds examined in the sensitivity analysis.

To estimate uncertainty, a dataset-level cluster bootstrap was performed because comparisons within each dataset shared the same reference set. For each of 2,000 bootstrap replicates, complete datasets were sampled with replacement, and suggestion rates were recalculated from the combined counts. The 2.5$^{th}$ and 97.5$^{th}$ percentiles of the bootstrap estimates defined the 95% confidence intervals. Bootstrap sampling used a separate random seed, and confidence intervals were not reported when no incorrect suggestions were observed.

Bootstrap stability was assessed using five additional random seeds with 2,000 resamples each and one additional seed with 10,000 resamples. Confidence interval bounds were compared with those from the original analysis using the same evaluation datasets.

Suggestion rates at the selected thresholds and at HYphlow's fixed thershold in the independent evaluation datasets are given in Supplementary Table S3. Reference names sharing a second word, such as "virus" or "bacterium", are counted in Supplementary Table S4.

**Supplementary Table S3. Suggestions at each threshold in the independent evaluation datasets Each group was evaluated with 1,000 datasets generated independently of threshold selection (3,000,000 spelling-variant and 10,000 absent-taxon comparisons).** Corr., the variant's own reference name was suggested; Incorr., a different reference name was suggested; None, no reference name reached the threshold. For absent taxa every suggestion is correct, and the remainder received no suggestion. A few spelling variants that exactly matched a different reference name are counted in no column. 95% CI from 2,000 dataset-level bootstrap replicates; no interval is given for zero events. Thresholds other than 67 were reviewed after the evaluation results were known. [a]Threshold selected in Eukaryota by the predefined rule and applied to all groups. [b]HYphlow's fixed threshold.

| Group | Threshold | Spelling variants | | | Absent tax |
|---|---|---|---|---|---|
| | | Corr. (%) | Incorr. (%) | None (%) | Incorr. (%) (95% CI) |
| Eukaryota | 67[a] | 99.7 | 0.0001 | 0.3 | 1.10 (0.91-1.29 |
| | 80[b] | 91.7 | <0.0001 | 8.3 | 0.05 (0.01-0.10) |
| | 82 | 85.2 | <0.0001 | 14.8 | 0.03 (0.00-0.07) |
| | 85 | 73.2 | <0.0001 | 26.8 | 0.02 (0.00-0.05) |
| | 88 | 57.7 | 0 | 42.3 | 0 |
| | 89 | 50.9 | 0 | 49.1 | 0 |
| | 90 | 47.2 | 0 | 52.8 | 0 |
| Fungi | 67[a] | 99.9 | 0.0005 | 0.1 | 5.77 (5.33-6.22) |
| | 80[b] | 94.5 | 0.0003 | 5.5 | 0.53 (0.40-0.70) |
| | 82 | 89.3 | 0.0001 | 10.7 | 0.28 (0.17-0.40) |
| | 85 | 78.6 | 0.0001 | 21.4 | 0.08 (0.03-0.14) |
| | 88 | 62.8 | <0.0001 | 37.2 | 0.01 (0.00-0.04) |
| | 89 | 55.9 | 0 | 44.1 | 0 |
| | 90 | 51.6 | 0 | 48.4 | 0 |
| Bacteria | 67[a] | 99.9 | 0.0008 | 0.1 | 15.01 (14.31-15.75) |
| | 80[b] | 96.4 | 0.0005 | 3.6 | 1.66 (1.40-1.91) |
| | 82 | 92.4 | 0.0003 | 7.6 | 0.81 (0.64-0.99) |
| | 85 | 83.3 | 0.0003 | 16.7 | 0.34 (0.23-0.46) |
| | 88 | 67.9 | 0.0001 | 32.1 | 0.15 (0.08-0.23) |
| | 89 | 60.6 | <0.0001 | 39.4 | 0.06 (0.02-0.11) |
| | 90 | 55.9 | <0.0001 | 44.1 | 0.04 (0.01-0.08) |
| Virus | 67[a] | 99.2 | 0.1879 | 0.7 | 26.15 (25.28-27.01) |
| | 80[b] | 90.1 | 0.0707 | 9.8 | 8.42 (7.85-8.98) |
| | 82 | 84.0 | 0.0428 | 16.0 | 4.42 (4.01-4.84) |
| | 85 | 73.5 | 0.0234 | 26.5 | 2.64 (2.32-2.99) |
| | 88 | 59.7 | 0.0142 | 40.3 | 0.82 (0.65-1.01) |
| | 89 | 52.6 | 0.0070 | 47.4 | 0.34 (0.23-0.46) |
| | 90 | 48.4 | 0.0061 | 51.5 | 0.32 (0.21-0.44) |

**Supplementary Table S4.** Reference names sharing a second word. Eligible names of each group were countered by their second word. Incorrect suggestions with score ≥85 are counts of spelling variants and absent taxa combined in the independent evaluation datasets (3,000,000 spelling-variant and 10,000 absent-taxon comparisons per group). Fungi is a subset of Eukaryota.

| Group | Eligible names | name "virus", n (%) | name "bacterium", n (%) | Incorrect suggestions with score ≥85, n |
|---|---|---|---|---|
| Eukaryota | 769,353 | 0 | 0 | 3 |
| Fungi | 68,242 | 0 | 0 | 11 |

| | | | | |
|---|---|---|---|---|
| Bacteria | 27,907 | 0 | 1,079 (3.9) | 42 |
| Viruses | 14,997 | 3,465 (23.1) | 0 | 967 |

### Supplementary Text S2. Evaluation of tree annotation on simulated phylogenies with known ancestral states

Ancestral states are generally unknown in empirical data, so the accuracy of tree annotation cannot be assessed directly. We therefore simulated phylogenies and binary traits with known ancestral states and used them as references. We first assessed whether HYphlow's MP and ML reconstructions agreed with established R packages and correctly recovered the simulated ancestral states. We then compared HYphlow's three foreground assignment methods and evaluated the default parameters of the consensus method.

#### *S2.1. Simulated trees and trait with known ancestral states*

Tip labels were drawn from a pool of 300 fictional binomial names in **Genus_epithet** format, generated from romanized Korean words using a fixed random seed. The names were not checked against taxonomic databases and carried no taxonomic or phylogenetic information.

Trees were generated with 8, 16, 32, 64, 128 and 256 tips, using three topologies and two branch-length conditions, with 10 replicates per combination (360 trees in total). For each tree size and replicate, taxa were sampled without replacement from the name pool and randomly assigned to tips. The same taxon set was used across all three topologies and both branch-length conditions, allowing comparisons using the same taxa. Balanced trees were generated by separating one taxon at each successive split (Fischer et al. 2023). These two topologies represent the extremes of tree balance measured by the Colless and Sackin indices (Sackin 1972; Colless 1982; Fischer et al. 2023). Yule topologies were generated by repeatedly selecting a tip uniformly at random and splitting it (Harding 1971). Yule branching times were not retained, so the same branch-length assignment procedure was used for all three topologies.

Because balanced and caterpillar topologies are fixed for each tree size, their replicates differed only in tip labels and branch lengths, and under the equal branch-length condition, only in tip labels. These replicates were therefore not topologically independent, but each received independently simulated traits. Separate random seeds for taxon sampling, topology generation, tip assignment and branch-length generation were derived from a fixed base seed, the simulation condition and the replicate number. The seeds were recorded for each tree. Random numbers in all Python analyses were generated using the Mersenne Twister generator (Python v3.12.12).

Branch lengths were either equal across all branches within a tree (equal condition) or drawn from an exponential distribution with rate 1 (variable condition). Both conditions were rescaled to a mean root-to-tip distance of 1, calculated by weighting each branch length by the proportion of tips descending from that branch and summing the weighted lengths. Total tree length was not standardized and varied among topologies (**Supplementary Table S5**). A minimum branch length of $10^{-4}$ was imposed, because zero-length branches constrain the states at both ends of a branch to be identical in both trait simulation and likelihood-based reconstruction. Branches below this minimum were raised to $10^{-4}$ , and the remaining branch lengths were adjusted to preserve the mean root-to-tip distance. Branch lengths were rounded to 10 significant digits, and the rounded values were used in all subsequent analyses.

**Supplementary Table S5. Characteristics of the simulated trees.** Total tree length is the sum of all branch lengths, averaged across 10 replicates per condition. Root-to-tip ranges indicate the minimum and maximum distances across replicates. Branches raised to minimum is the total number of branches increased to $10^{-4}$ under the variable branch-length condition, out of 10 * (2n-2) branches per condition, where n is the number of tips. All trees were scaled to a mean root-to-tip distance of 1 and passed structural validation.

| Topology | Tips | Total length | | Root-to-tip range | | Branches raised to minimum |
|---|---|---|---|---|---|---|
| | | Equal | Variable | Equal | Variable | |
| Balanced | 8 | 4.67 | 4.52 | 1.00-1.00 | 0.32-2.48 | 0 |
| | 16 | 7.50 | 7.71 | 1.00-1.00 | 0.23-2.20 | 0 |
| | 32 | 12.40 | 12.82 | 1.00-1.00 | 0.11-2.48 | 1 |
| | 64 | 21.00 | 20.12 | 1.00-1.00 | 0.22-2.84 | 0 |
| | 128 | 36.29 | 33.23 | 1.00-1.00 | 0.22-2.49 | 0 |
| | 256 | 63.75 | 65.86 | 1.00-1.00 | 0.25-2.69 | 7 |
| Caterpillar | 8 | 3.20 | 3.46 | 0.23-1.60 | 0.0015-2.11 | 0 |
| | 16 | 3.56 | 3.80 | 0.12-1.78 | 0.0063-2.44 | 1 |
| | 32 | 3.77 | 4.04 | 0.061-1.88 | 0.0026-2.16 | 0 |
| | 64 | 3.88 | 3.99 | 0.031-1.94 | 0.0014-2.29 | 5 |
| | 128 | 3.94 | 4.10 | 0.016-1.97 | 0.0017-2.28 | 15 |
| | 256 | 3.97 | 3.95 | 0.0078-1.98 | 0.00074-2.06 | 64 |
| Yule | 8 | 4.21 | 4.20 | 0.28-1.60 | 0.15-2.51 | 0 |
| | 16 | 6.35 | 6.28 | 0.20-1.86 | 0.040-2.22 | 0 |
| | 32 | 9.83 | 10.30 | 0.28-1.75 | 0.042-2.56 | 0 |
| | 64 | 16.28 | 17.40 | 0.12-1.97 | 0.083-2.41 | 0 |
| | 128 | 29.58 | 31.25 | 0.22-1.67 | 0.053-2.37 | 4 |
| | 256 | 49.46 | 47.70 | 0.19-1.85 | 0.061-2.63 | 7 |

All 360 trees were validated using ETE3 v3.1.3 and DendroPy v5.1.0, with code independent of the tree generation script. The 29 validation criteria included 23 applied to individual trees and six applied to the complete tree set. These criteria covered tree structure, tip labels, minimum branch length, mean root-to-tip distance (tolerance $10^{-8}$), agreement between ETE3 and DendroPy, and descendant tip set used to match nodes between programs. Balanced and caterpillar trees were also checked against their expected Colless, Sackin and cherry counts.

For Yule topologies, mean cherry counts and Sackin indices were compared with their theoretical expectations of $n/3$ (McKenzie and Steel 2000) and $2n(H_n - 1)$ (Kirkpatrick and Slatkin 1993), respectively, where $n$ is the number of tips and $H_n$ is the $n$th harmonic number. Standardized deviations ($z$) were calculated using the standard error across the 10 equal-length Yule trees at each size. Variable-length Yule trees were not included separately because they shared their topologies with the equal-length trees. The flagging threshold was set at $|z| > 3$ before analysis. No Yule statistic exceeded this threshold (maximum $|z| = 2.45$).

The coefficient of variation of variable branch lengths ranged from 0.84 to 1.05 compared with a reference value of 1 for an exponential distribution. This was recorded as a descriptive check rather than a validation criterion because the minimum branch length and root-to-tip scaling could alter the coefficient of variation.

Binary traits, with state 1 denoting the foreground and state 0 the background, were simulated under the five scenarios in **Table 3**. SYM and ARD followed continuous-time Markov models of discrete-character evolution (Pagel 1994; Lewis 2001). In SYM, gain ($0 \rightarrow 1$)and loss ($1 \rightarrow 0$) rates were equal ($q_{01} = q_{10} = \mu$). In ARD, the gain rate was four times the loss rate ($q_{01} = 1.6\mu$ , $q_{10} = 0.4\mu$), keeping the sum of rates equal to that in SYM. RV followed SYM, except that each branch's transition rate was multiplied by a factor drawn from a gamma distribution with shape 1 and mean 1 (equivalent to an exponential distribution). These factors were rescaled to a weighted mean of 1 based on each branch's contribution to the mean root-to-tip distance, preserving a mean transition rate of $\mu$ across root-to-tip paths. For all three scenarios, the root state was assigned as 0 or 1 with equal probability, matching HYphlow's root prior rather than the stationary distribution of ARD. Trait histories were simulated in continuous time from the root, with ancestral states and the position and direction of each transition recorded.

For SG, the root state was fixed at 0, and a single gain was placed at a position drawn uniformly along the total tree length, so that the gain branch was selected with probability proportional to its length. All nodes descending from the gain were assigned state 1, and no losses occurred. For NS, tip states were assigned independently of phylogeny with a foreground probability of 0.25 or 0.5 (Maddison and Slatkin 1991). No ancestral states were simulated for NS datasets. An additional rate of $\mu = 0.02$ was initially simulated but excluded before ancestral state reconstruction because most datasets at this rate had uniform tip states. Separate random seeds for root states, trait histories, gain positions, rate multipliers and tip states were derived from each dataset identifier and recorded.

All 5,400 trait datasets were validated against 20 criteria using code independent of the trait simulation script. Sixteen criteria were applied to individual datasets and four to the complete datasets. These criteria covered trait tables read by HYphlow, agreement between observed and recorded tip states, scenario-specific transition rates, rescaling of RV multipliers (tolerance $10^{-9}$), the single-gain structure of SG histories and the absence of simulated ancestral states in NS datasets. Replaying the recorded transition from the root reproduced all recorded ancestral states.

Simulated traits were also assessed against theoretical expectations using 46 diagnostics pooled across trees. These included root-state frequencies, branches with different start and end states, transition counts relative to time spent in each state, variation in transition counts among branches, SG gain positions, RV multipliers before rescaling, and NS tip frequencies and sister-tip agreement. Flagging thresholds were specified before analysis at $|z| > 3$ for standardized deviations and $p < 0.001$ for other tests. No diagnostic exceeded these thresholds (maximum $|z| = 2.61$).

In total, 5,400 datasets were simulated (360 trees * 15 trait conditions), of which 4,680 had known ancestral states. Datasets with uniform tip states were retained and flagged rather than regenerated but excluded from reconstruction comparisons (**Supplementary Table S6**). This left 4,810 datasets for agreement analyses, including 713 NS datasets, and 4,097 datasets with known ancestral states for accuracy analyses.

**Supplementary Table S6. Datasets with uniform tip states by evolutionary scenarios, transition rates and tree size**. Values indicate the number of datasets with uniform tip states out of 60 per cell (3 topologies * 2 branch-length conditions * 10 replicates). Of the 4,680 datasets with known ancestral states, 583 had uniform tip states and were excluded from reconstruction comparisons. Uniform tip states were most frequent at $\mu = 0.1$ and in smaller trees, where fewer transitions were expected. Under ARD, they also occurred at higher transition rates, consistent with the stationary probability of 0.8 for state 1. NS datasets are not shown because ancestral states were not simulated for them. Seven NS datasets with uniform tip states were also excluded from agreement analyses.

| Scenario | μ | 8 | 16 | 32 | 64 | 128 | 256 | Total (of 360) |
|---|---|---|---|---|---|---|---|---|
| SYM | 0.1 | 40 | 36 | 25 | 19 | 16 | 10 | 146 |
| | 0.5 | 9 | 7 | 3 | 4 | 1 | 1 | 25 |
| | 2.5 | 1 | 0 | 0 | 0 | 0 | 0 | 1 |
| | 10 | 1 | 0 | 0 | 0 | 0 | 0 | 1 |
| ARD | 0.1 | 38 | 36 | 29 | 26 | 17 | 17 | 163 |
| | 0.5 | 17 | 13 | 5 | 3 | 0 | 5 | 43 |
| | 2.5 | 9 | 4 | 0 | 0 | 0 | 0 | 13 |
| | 10 | 11 | 1 | 0 | 0 | 0 | 0 | 12 |
| RV | 0.1 | 41 | 34 | 25 | 15 | 16 | 17 | 148 |
| | 0.5 | 10 | 7 | 3 | 5 | 4 | 0 | 29 |
| | 2.5 | 1 | 1 | 0 | 0 | 0 | 0 | 2 |
| | 10 | 0 | 0 | 0 | 0 | 0 | 0 | 0 |
| SG | — | 0 | 0 | 0 | 0 | 0 | 0 | 0 |
| **Total** | | | | | | | | **583** |

#### *S2.2 Agreement of HYphlow's parsimony and likelihood reconstructions with established R packages*

The maximum parsimony (MP) and maximum likelihood (ML) reconstructions produced by HYphlow were evaluated for agreement with those produced by phangorn v2.12.1 (Schliep 2011) and castor v1.8.7 (Louca and Doebeli 2018), respectively, both implemented in R v4.3.3 (R Core Team 2024). HYphlow and the corresponding R packages received the same rooted, bifurcating tree and observed tip states for each dataset.

The R packages were selected because they implement the same reconstruction methods and models as HYphlow, allowing comparisons under the same reconstruction criteria. In phangorn, MP reconstruction used `ancestral.pars` with `type = "MPR"` to identify states occurring in at least one most parsimonious reconstruction (MPR), consistent with HYphlow's Fitch parsimony. In castor, ML reconstruction used `asr_mk_model` with the same two-state equal-rates model (`rate_model = "ER"`) and flat root prior (`root_prior = "flat"`) as HYphlow. Marginal ancestral state probabilities at each node, given all observed tip states, were calculated using `reroot = TRUE`. When the transition rate was estimated, castor used five random starting points (`Ntrials = 5`) with a fixed random seed. All other arguments were left at their package defaults. Reconstructions were compared at the root and all internal nodes of each tree. Tips were excluded because their states were observed rather than reconstructed. Because node numbering differs between programs, nodes were matched using their descendant taxa.

For MP, HYphlow and phangorn returned identical MPR state sets at all 417,886 nodes across the 4,810 datasets. Agreement was complete across all simulation scenarios, transition rates, tree sizes, topologies and branch-length conditions, including NS datasets. With no disagreements, the upper 95% confidence bound for the proportion of datasets with any disagreement was 0.0006 overall (3/4,810, using the rule of three (Hanley and Lippman-Hand 1983)) and at most 0.015 within individual simulation conditions (**Supplementary Table S7**).

**Supplementary Table S7. Agreement between HYphlow and phangorn MPR state sets at the root and internal nodes.** The upper 95% bound is one0sided bound for the proportion of datasets with any disagreement, from the rule of three (3/*n*). $\mu$, simulated transition rate; $\rho$, NS foreground probability.

| Group | Datasets | Nodes | Nodes in agreement (%) | Upper 95% bound |
|---|---|---|---|---|
| **All** | 4,810 | 417,886 | 100 | 0.0006 |
| ***Simulation condition*** | | | | |
| SYM ($\mu$=0.1) | 214 | 22,506 | 100 | 0.0140 |
| SYM ($\mu$=0.5) | 335 | 28,985 | 100 | 0.0090 |
| SYM ($\mu$=2.5) | 359 | 29,873 | 100 | 0.0084 |
| SYM ($\mu$=10) | 359 | 29,873 | 100 | 0.0084 |
| ARD ($\mu$=0.1) | 197 | 20,043 | 100 | 0.0152 |
| ARD ($\mu$=0.5) | 317 | 27,947 | 100 | 0.0095 |
| ARD ($\mu$=2.5) | 347 | 29,757 | 100 | 0.0086 |
| ARD ($\mu$=10) | 348 | 29,788 | 100 | 0.0086 |
| RV ($\mu$=0.1) | 212 | 20,996 | 100 | 0.0142 |
| RV ($\mu$=0.5) | 331 | 28,789 | 100 | 0.0091 |
| RV ($\mu$=2.5) | 358 | 29,858 | 100 | 0.0084 |
| RV ($\mu$=10) | 360 | 29,880 | 100 | 0.0083 |
| SG | 360 | 29,880 | 100 | 0.0083 |
| NS ($\rho$=0.25) | 353 | 29,831 | 100 | 0.0085 |
| NS ($\rho$=0.5) | 360 | 29,880 | 100 | 0.0083 |
| ***Tree size (tips)*** | | | | |
| 8 | 715 | 5,005 | 100 | 0.0042 |
| 16 | 761 | 11,415 | 100 | 0.0039 |

| | | | | |
|---|---|---|---|---|
| 32 | 810 | 25,110 | 100 | 0.0037 |
| 64 | 828 | 52,164 | 100 | 0.0036 |
| 128 | 846 | 107,442 | 100 | 0.0035 |
| 256 | 850 | 216,750 | 100 | 0.0035 |
| ***Topology*** | | | | |
| Balanced | 1,667 | 145,861 | 100 | 0.0018 |
| Caterpillar | 1,474 | 125,422 | 100 | 0.0020 |
| Yule | 1,669 | 146,603 | 100 | 0.0018 |
| ***Branch lengths*** | | | | |
| Equal | 2,425 | 210,415 | 100 | 0.0012 |
| Variable | 2,385 | 207,471 | 100 | 0.0013 |

For ML, reconstructions from HYphlow and castor were compared using two approaches. In the fixed-rate comparison, both implementations received the transition rate used to simulate each SYM dataset (`transition_matrix` in castor). This allowed marginal ancestral probabilities to be compared without differences in rate estimation. The fixed-rate comparison was limited to SYM because it was the only simulation scenario with a single symmetric transition rate matching the reconstruction model. In the estimated-rate comparison, each implementation estimated its own transition rate, as in a standard analysis.

Marginal foreground probabilities were compared at each node, with absolute differences $\leq 10^{-4}$ considered probability agreement. Nodes with probabilities greater than 0.5 were assigned to the foreground state, and the remaining nodes to the background state. Nodes with probabilities within $10^{-4}$ of 0.5 in either implementation were classified as ties and excluded from state agreement calculations. These nodes were recorded separately because small numerical differences could change their assigned states.

In the fixed-rate comparison, HYphlow and castor assigned the same state to 108,700 of 108,701 nodes across 1,267 SYM datasets, excluding 2,536 nodes classified as ties. The only disagreement occurred near the 0.5 threshold, with marginal foreground probabilities of 0.5007 in HYphlow and 0.4993 in castor. Marginal probabilities differed by no more than $10^{-4}$ at 107,277 of 111,237 nodes (96.4%), with a median absolute difference of $4.9 * 10^{-7}$ and a maximum of $6.3 * 10^{-3}$.

The larger differences arose from the numerical approximation castor uses to calculate branch transition probabilities. For the 541 datasets with differences greater than $10^{-4}$, castor's marginal probabilities were recalculated using exact transition probabilities. The maximum difference from HYphlow decreased to $6.1 * 10^{-6}$, and no node exceeded $10^{-4}$. This recalculation was used only to identify the source of the differences. The primary comparisons used castor's default settings.

In the estimated-rate comparison, HYphlow and castor assigned the same state to 385,783 of 385,847 nodes (99.98%) across 4,810 datasets, excluding 32,039 nodes classified as ties. The 64 disagreements occurred in 15 datasets, including 12 NS datasets. Marginal probabilities differed by no more than $10^{-4}$ at 381,711 of 417,886 (91.3%), with a median absolute difference of $7.3 * 10^{-9}$. Probability agreement was lowest for NS datasets (74.8%) and ranged from 93.3% to 99.6% across the remaining simulation scenarios (**Supplementary Table S8**). Estimated transition rates also differed between the two implementations. HYphlow searches rates up to 100, whereas castor's estimates are unbounded. In 510 of the 4,810 datasets (10.6%), including 224 of the 713 NS datasets (31.4%), HYphlow's estimate reached this upper limit, while castor's estimates for these datasets ranged from 2.9 to 17,096 (median 809). In the remaining datasets, the two estimates differed by less than 1% in 85.5% of datasets.

**Supplementary Table S8. Agreement between HYphlow and castor ML reconstructions.** Nodes include the root and all internal nodes of each tree. In the estimated-rate comparison, each implementation estimated its own transition rate. In the fixed-rate comparison, both implementations used the simulated transition rate. This comparison was limited to SYM datasets, the only scenario with a single symmetric rate matching the reconstruction model (— indicates not applicable). State agreement is the percentage of nodes assigned the same state by both implementations, excluding ties. Ties were defined as nodes where either implementation's marginal foreground probability was within $10^{-4}$ of 0.5. Probability agreement is the percentage of nodes whose marginal foreground probabilities differed by no more than $10^{-4}$ between implementations. $\mu$ denotes the simulated transition rate, and $\rho$ denotes the foreground proportion in NS datasets.

| Group | Datasets | Nodes | Estimated rate: State (%) | Ties | Probability (%) | Fixed rate: State (%) | Ties | Probability (%) |
|---|---|---|---|---|---|---|---|---|
| **All** | 4,810 | 417,886 | 99.98 | 32,039 | 91.3 | — | — | — |
| ***Simulation condition*** | | | | | | | | |
| **SYM (all rates)** | **1,267** | **111,237** | **100** | **7,600** | **93** | **>99.99** | **2,536** | **96.4** |

| | | | | | | | | |
|---|---|---|---|---|---|---|---|---|
| SYM ($\mu$=0.1) | 214 | 22,506 | 100 | 13 | 99.3 | 100 | 7 | 98.0 |
| SYM ($\mu$=0.5) | 335 | 28,985 | 100 | 117 | 96.2 | >99.99 | 2 | 96.0 |
| SYM ($\mu$=2.5) | 359 | 29,873 | 100 | 1,285 | 90.1 | 100 | 34 | 94.5 |
| SYM ($\mu$=10) | 359 | 29,873 | 100 | 6,185 | 89.2 | 100 | 2,493 | 97.6 |
| **ARD (all rates)** | **1,209** | **107,535** | **>99.99** | **798** | **93.6** | **—** | **—** | **—** |
| ARD ($\mu$=0.1) | 197 | 20,043 | 100 | 4 | 99.3 | — | — | — |
| ARD ($\mu$=0.5) | 317 | 27,947 | >99.99 | 66 | 96.5 | — | — | — |
| ARD ($\mu$=2.5) | 347 | 29,757 | 100 | 278 | 92.5 | — | — | — |
| ARD ($\mu$=10) | 348 | 29,788 | >99.99 | 450 | 88.2 | — | — | — |
| **RV (all rates)** | **1,261** | **109,523** | **>99.99** | **4,262** | **93.8** | **—** | **—** | **—** |
| RV ($\mu$=0.1) | 212 | 20,996 | 100 | 1 | 99.1 | — | — | — |
| RV ($\mu$=0.5) | 331 | 28,789 | 100 | 52 | 96.4 | — | — | — |
| RV ($\mu$=2.5) | 358 | 29,858 | >99.99 | 626 | 90.7 | — | — | — |
| RV ($\mu$=10) | 360 | 29,880 | 100 | 3,583 | 90.7 | — | — | — |
| SG | 360 | 29,880 | 100 | 3 | 99.6 | — | — | — |
| **NS (all)** | **713** | **59,711** | **99.85** | **19,376** | **74.8** | — | — | — |
| NS ($\rho$=0.25) | 353 | 29,831 | 99.94 | 2,825 | 80.1 | — | — | — |
| NS ($\rho$=0.5) | 360 | 29,880 | 99.67 | 16,551 | 69.6 | — | — | — |
| ***Tree size (tip)*** | | | | | | | | |
| 8 | 715 | 5,005 | 100 | 1,445 | 91.3 | — | — | — |
| 16 | 761 | 11,415 | 100 | 2,147 | 92.0 | — | — | — |
| 32 | 810 | 25,110 | 100 | 2,689 | 91.3 | — | — | — |
| 64 | 828 | 52,164 | 100 | 4,929 | 91.4 | — | — | — |
| 128 | 846 | 107,442 | >99.99 | 7,072 | 92.0 | — | — | — |
| 256 | 850 | 216,750 | 99.97 | 13,757 | 91.0 | — | — | — |
| ***Topology*** | | | | | | | | |
| Balanced | 1,667 | 145,861 | >99.99 | 13,831 | 92.7 | — | — | — |
| Caterpillar | 1,474 | 125,422 | 99.95 | 5,582 | 90.0 | — | — | — |
| Yule | 1,669 | 146,603 | >99.99 | 12,626 | 91.2 | — | — | — |
| ***Branch lengths*** | | | | | | | | |
| Equal | 2,425 | 210,415 | 100 | 18,144 | 98.6 | — | — | — |
| Variable | 2,385 | 207,471 | 99.97 | 13,895 | 84.0 | — | — | — |

#### *S2.3 Accuracy of reconstructed ancestral states*

The MP and ML reconstructions produced by HYphlow, phangorn and castor were evaluated for accuracy against simulated ancestral states. NS datasets were excluded because ancestral states were not simulated for them. Simulated ancestral states were stored separately and were not available to any implementation during reconstruction. No reconstruction script read the file containing them. All implementations were evaluated on the same 358,175 root and internal nodes.

For MP, nodes whose MPR state sets contained both states were classified as ambiguous. Accuracy was calculated as the proportion of unambiguous nodes whose single MPR state matched the simulated state. The proportion of ambiguous nodes was reported separately. For ML, each implementation estimated its own transition rate and nodes with marginal foreground probabilities greater than 0.5 were assigned to the foreground state. Accuracy was calculated as the proportion of all nodes assigned the simulated state. Unlike in the agreement comparison, nodes with probabilities close to 0.5 were not excluded. Marginal probabilities were also evaluated using the Brier score (Brier 1950; Gneiting and Raftery 2007):

$$BS = \frac{1}{N}\sum_{i=1}^{N}(p_i - y_i)^2$$

where $p_i$ is the marginal foreground probability at node $i$, $y_i$ is the simulated state (0 or 1), and $N$ is the number of evaluated nodes. Lower scores indicate that the predicted probabilities are closer to the simulated states.

ARD, RV and SG datasets were simulated under models that differed from the symmetric two-state model used for ML reconstruction. These datasets were included to evaluate reconstruction accuracy when the assumed model differed from the simulated evolutionary process. Overall accuracy was calculated by pooling node counts across all datasets. Accuracy was also calculated separately by simulation scenario, tree topology, branch-length condition and tree size. Because nodes within a tree are not independent, 95% confidence intervals (CI) were calculated using a cluster bootstrap. Each cluster contained all datasets from the same tree size and replicate, giving 60 clusters in total. These datasets shared the same taxon set, and trees differing only in branch-length condition shared the same topology.

For each of 2,000 bootstrap replicates, 10 clusters were drawn with replacement within each tree size, preserving the original distribution of tree sizes. Accuracy, ambiguity proportions and Brier scores were recalculated from the pooled node counts for each replicate. Differences in accuracy and Brier score were calculated as HYphlow minus castor within the same bootstrap replicates. The 95% CI was defined by the 2.5th and 97.5th percentiles of the 2,000 bootstrap values for each measure. A random seed was recorded to allow the bootstrap to be reproduced.

MP reconstructions were ambiguous at 5.8% of the 358,175 nodes (95% CI 5.6-6.0%). Among unambiguous nodes, 91.9% matched the simulated ancestral state (95% CI 91.8-92.1%). As tree size increased from 8 to 256 tips, the proportion of ambiguous nodes decreased from 14.0% to 4.9%, while accuracy increased from 81.4% to 93.2%. Accuracy was highest for caterpillar trees (98.0%) and lowest for balanced trees (88.5%). Accuracy declined with increasing simulated transition rates. At $\mu = 0.1$, accuracy exceeded 99.9%, but at $\mu = 10$, it decreased to 71.2% for SYM, 85.0% for ARD and 76.6% for RV. In SG datasets, all ambiguous nodes matched the simulated sate, and 0.05% of nodes were ambiguous (**Supplementary Table S9**).

**Supplementary Table S9. Accuracy of MP ancestral state reconstructions at the root and internal nodes of the 4,097 datasets with known ancestral states.** HYphlow and phangorn returned identical MPR state sets, so one value is shown for both. Accuracy is the percentage of unambiguous nodes whose MPR state matched the simulated state. 95% CIs are from 2,000 cluster bootstrap replicates. *μ,* simulated transition rate.

| Group | Nodes | Ambiguous (%) [95% CI] | Accuracy (%) [95% CI] |
|---|---|---|---|
| **All** | 358,175 | 5.78 [5.59, 5.97] | 91.95 [91.79, 92.12] |
| ***Simulation condition*** | | | |
| SYM (*μ*=0.1) | 22,506 | 0.23 [0.14, 0.33] | 99.95 [99.92, 99.98] |
| SYM (*μ*=0.5) | 28,985 | 2.04 [1.83, 2.28] | 98.89 [98.64, 99.09] |
| SYM (*μ*=2.5) | 29,873 | 12.23 [11.19, 13.30] | 86.72 [85.93, 87.41] |
| SYM (*μ*=10) | 29,873 | 18.59 [17.27, 19.92] | 71.15 [70.51, 71.78] |
| ARD (*μ*=0.1) | 20,043 | 0.22 [0.14, 0.31] | 99.97 [99.95, 99.99] |
| ARD (*μ*=0.5) | 27,947 | 1.97 [1.64, 2.31] | 98.84 [98.61, 99.05] |
| ARD (*μ*=2.5) | 29,757 | 3.34 [2.89, 3.81] | 90.07 [89.36, 90.71] |
| ARD (*μ*=10) | 29,788 | 3.46 [3.03, 3.89] | 85.01 [84.56, 85.46] |
| RV (*μ*=0.1) | 20,996 | 0.21 [0.13, 0.29] | 99.91 [99.87, 99.95] |
| RV (*μ*=0.5) | 28,789 | 1.76 [1.47, 2.05] | 98.99 [98.83, 99.14] |
| RV (*μ*=2.5) | 29,858 | 10.14 [9.40, 10.90] | 90.10 [89.44, 90.72] |
| RV (*μ*=10) | 29,880 | 15.48 [14.38, 16.59] | 76.64 [75.93, 77.39] |
| SG | 29,880 | 0.05 [0.03, 0.08] | 100.00 [100.00, 100.00] |
| ***Tree size (tips)*** | | | |
| 8 | 4,214 | 13.98 [11.95, 16.31] | 81.41 [79.79, 82.80] |
| 16 | 9,615 | 10.70 [9.46, 12.10] | 84.73 [83.89, 85.62] |
| 32 | 21,390 | 7.39 [6.63, 8.10] | 88.41 [87.71, 89.08] |
| 64 | 44,604 | 6.62 [6.17, 7.13] | 90.39 [90.14, 90.63] |
| 128 | 92,202 | 5.95 [5.48, 6.48] | 92.11 [91.74, 92.52] |
| 256 | 186,150 | 4.86 [4.66, 5.06] | 93.20 [92.96, 93.41] |

| | | | |
|---|---|---|---|
| ***Topology*** | | | |
| Balanced | 125,948 | 7.53 [7.18, 7.90] | 88.48 [88.17, 88.78] |
| Caterpillar | 105,530 | 2.26 [2.15, 2.39] | 98.03 [97.91, 98.14] |
| Yule | 126,697 | 6.96 [6.56, 7.35] | 90.05 [89.73, 90.39] |
| ***Branch lengths*** | | | |
| Equal | 180,563 | 5.99 [5.70, 6.31] | 91.25 [91.01, 91.49] |
| Variable | 177,612 | 5.56 [5.29, 5.83] | 92.65 [92.46, 92.83] |

ML accuracy was 90.6% (95% CI 90.5-90.8%) for HYphlow and 90.4% (95% CI 90.2-90.6%) for castor, with Brier scores of 0.0647 and 0.0648, respectively. Accuracy differed by 0.22 percentage points between the two implementations (HYphlow - castor, 95% CI 0.18-0.27), and Brier scores by -0.00007 (95% CI -0.00011 to -0.00005). Accuracy increased with tree size in both implementations, from 75.3% for HYphlow and 74.1% for castor at 8 tips to 92.1% and 92.0%, respectively, at 256 tips. As observed for MP, accuracy declined with increasing simulated transition rates (**Supplementary Table S10**).

**Supplementary Table S10. Accuracy of ML ancestral state reconstructions with transition rate estimated by each implementation.** Accuracy is the percentage of nodes assigned the simulated state. Difference is HYphlow minus castor accuracy in percentage points, with 95% CIs . Lower Brier scores indicate probabilities closer to the simulated states.

| Group | Nodes | Accuracy (%): HYphlow | Accuracy (%): castor | Difference (pp) [95% CI] | Brier: HYphlow | Brier: castor |
|---|---|---|---|---|---|---|
| **All** | 358,175 | 90.61 | 90.39 | 0.22 [0.18, 0.27] | 0.0647 | 0.0648 |
| ***Simulation condition*** | | | | | | |
| SYM ($\mu$=0.1) | 22,506 | 99.88 | 99.88 | 0.00 [-0.01, 0.00] | 0.0011 | 0.0011 |
| SYM ($\mu$=0.5) | 28,985 | 98.15 | 98.14 | 0.01 [0.00, 0.04] | 0.0131 | 0.0131 |
| SYM ($\mu$=2.5) | 29,873 | 84.93 | 84.52 | 0.41 [0.22, 0.63] | 0.1023 | 0.1024 |
| SYM ($\mu$=10) | 29,873 | 70.12 | 69.36 | 0.76 [0.47, 1.08] | 0.1775 | 0.1778 |
| ARD ($\mu$=0.1) | 20,043 | 99.88 | 99.88 | 0.00 [0.00, 0.00] | 0.0008 | 0.0008 |
| ARD ($\mu$=0.5) | 27,947 | 98.38 | 98.38 | 0.00 [-0.01, 0.01] | 0.0130 | 0.0130 |
| ARD ($\mu$=2.5) | 29,757 | 90.29 | 90.10 | 0.19 [0.06, 0.36] | 0.0798 | 0.0799 |
| ARD ($\mu$=10) | 29,788 | 84.28 | 84.08 | 0.20 [0.06, 0.37] | 0.1339 | 0.1340 |
| RV ($\mu$=0.1) | 20,996 | 99.83 | 99.83 | 0.00 [0.00, 0.00] | 0.0013 | 0.0013 |
| RV ($\mu$=0.5) | 28,789 | 98.36 | 98.35 | 0.01 [0.00, 0.03] | 0.0122 | 0.0122 |
| RV ($\mu$=2.5) | 29,858 | 88.08 | 87.92 | 0.16 [0.07, 0.26] | 0.0842 | 0.0842 |
| RV ($\mu$=10) | 29,880 | 74.86 | 73.93 | 0.93 [0.56, 1.37] | 0.1601 | 0.1604 |
| SG | 29,880 | 99.98 | 99.98 | 0.00 [0.00, 0.00] | 0.0002 | 0.0002 |
| ***Tree size (tips)*** | | | | | | |
| 8 | 4,214 | 75.27 | 74.11 | 1.16 [0.69, 1.63] | 0.1549 | 0.1556 |
| 16 | 9,615 | 81.08 | 79.96 | 1.12 [0.68, 1.61] | 0.1278 | 0.1284 |
| 32 | 21,390 | 86.76 | 86.18 | 0.58 [0.37, 0.87] | 0.0945 | 0.0947 |
| 64 | 44,604 | 89.08 | 88.63 | 0.45 [0.30, 0.61] | 0.0757 | 0.0758 |
| 128 | 92,202 | 90.89 | 90.74 | 0.15 [0.08, 0.22] | 0.0627 | 0.0628 |

| | | | | | | |
|---|---|---|---|---|---|---|
| 256 | 186,150 | 92.12 | 92.03 | 0.10 [0.03, 0.16] | 0.0544 | 0.0544 |
| ***Topology*** | | | | | | |
| Balanced | 125,948 | 86.82 | 86.60 | 0.22 [0.14, 0.31] | 0.0886 | 0.0887 |
| Caterpillar | 105,530 | 97.39 | 97.32 | 0.07 [0.04, 0.09] | 0.0194 | 0.0195 |
| Yule | 126,697 | 88.74 | 88.38 | 0.36 [0.24, 0.48] | 0.0787 | 0.0789 |
| ***Branch lengths*** | | | | | | |
| Equal | 180,563 | 89.04 | 88.81 | 0.22 [0.16, 0.29] | 0.0739 | 0.0739 |
| Variable | 177,612 | 92.21 | 91.99 | 0.22 [0.16, 0.29] | 0.0554 | 0.0555 |

#### *S2.4 Comparison of HYphlow's foreground branch assignment methods*

Foreground branch assignments produced by HYphlow's MP, ML and consensus methods were compared with the simulated branch states in the 4,097 datasets with known ancestral states. Because phangorn and castor do not assign foreground branches, this comparison included only HYphlow.

#### *S2.5 Sensitivity of consensus assignments to the threshold and sample size*

HYphlow's consensus foreground assignment depends on two parameters: the sampled foreground proportion threshold (default 0.5) and the number of ancestral state samples (default 5,000). Consensus assignments were recalculated at thresholds from 0.3 to 0.9 in increments of 0.1, while the MP and ML reconstructions and sampled foreground proportions remained unchanged. At 0.5, the recalculated assignments matched HYphlow's original assignments for every branch. Increasing the threshold from 0.5 to 0.9 raised precision from 0.928 to 0.967 and reduced recall from 0.875 to 0.778. At 0.3 and 0.4, all 176,298 consensus-eligible branches were assigned to the foreground because every eligible branch had a sampled foreground proportion above 0.4. Increasing the threshold from 0.4 to 0.5 excluded 1,603 branches, of which 1,019 were simulated foreground branches.

The effect of the threshold was greater at higher transition rates. In SYM and RV datasets at $\mu = 2.5$ and 10, increasing the threshold from 0.5 to 0.9 reduced recall by 0.18-0.26 and increased precision by 0.06-0.19. In ARD datasets, recall decreased by 0.05 and 0.12 at these respective rates. At $\mu \leq 0.5$, recall decreased by at most 0.02 in SYM, ARD, and RV datasets.

To assess the effect of ancestral state sample size on consensus stability, sampling was repeated across the 4,097 datasets using 500, 1,000, 2,000, 5,000 and 10,000 samples, with 20 random seeds per dataset and sample size. MP and ML reconstructions and the consensus threshold of 0.5 remained unchanged. Before repeated sampling, each dataset was rerun with its recorded seed and 5,000 samples. The MP and ML reconstructions sampled foreground proportions and consensus assignments matched the original analyses in all 4,097 datasets.

Disagreement was calculated from Fleiss's (1971) observed agreement, which summarizes how consistently multiply raters assign items to categories. Here, each random seed was treated as a rater assigning each internal branch to the foreground or background:

$$D = \frac{\sum_{i=1}^{B} k_i(S - k_i)}{B \cdot S(S-1)/2}$$

where $k_i$ is the number of seeds assigning branch ($i$) to the foreground, $S$=20 is the number of seeds, and $B$ is the total number of internal branches across all datasets. Disagreement was therefore pooled across datasets. $D$ equals one minus Fleiss's observed agreement and represents the probability that two analyses with different seeds assign different states to a randomly chosen branch. A value of zero indicates that all seeds agreed on every branch. Observed agreement was used rather than Fleiss's kappa because the analysis assessed how often repeated assignments differed, rather than agreement adjusted for chance. A stability target of no more than 1% disagreement across all branches was specified before analysis.

As the number of ancestral state samples increases, the sampled foreground proportion of each eligible branch approaches its ML foreground probability. The limiting consensus assignment can therefore be determined without additional sampling. Because consensus-eligible branches have ML foreground probabilities above 0.5, every eligible branch is assigned to the foreground in the limit, and every ineligible branch to the background. Agreement with these limiting assignments increased from 99.38% at 500 samples to 99.54% at 5,000 samples and 99.57% at 10,000 samples.

Observed disagreement was also compared with the disagreement expected under binomial sampling. For an eligible branch with ML foreground probability $p$ and $n$ samples, the probability of a foreground assignment is $q = P(X > n/2)$, where $X$ follows a binomial distribution with $n$ trials and success probability $p$. The expected disagreement between two independent seeds is then $2q(1\text{-}q)$, with $q = 0$ for ineligible branches. Expected disagreement was averaged across all internal branches. Observed and expected disagreement differed by no more than

0.0012 percentage points at any sample size. This close agreement indicates that the observed variation across seeds was consistent with binomial sampling around the ML foreground probabilities.

Consensus assignments remained variable among branches with ML foreground probabilities close to 0.5. At 5,000 samples, 4,026 branches received different assignments across the 20 seeds. Of these, 63.7% had ML foreground probabilities within 0.001 of 0.5, 92.7% were within 0.01, and all were within 0.025. All 2,563 eligible branches with ML probabilities within 0.001 of 0.5 were unstable at every sample size, with mean disagreement near 0.50. Approximately 250,000 samples would be required for a foreground probability of 0.501 to differ from 0.5 by one standard error.

Because most internal branches had ML probabilities far from 0.5, disagreement was also calculated separately for consensus-eligible branches, giving 1.37%, 1.23%, 1.12%, 0.999% and 0.93% at 500, 1,000, 2,000, 5,000 and 10,000 samples, respectively. Across all internal branches, disagreement exceeded 1% at every sample size in SYM and RV datasets at $\mu = 10$ (2.6-3.5% and 1.7-2.8%, respectively) and in 16-tip trees (1.2-1.4%). These conditions had higher proportions of eligible branches with ML foreground probabilities between 0.5 and 0.6: 30% for SYM and 26% for RV at $\mu = 10$, and 9.6% for 16-tip trees, compared with 4.7% overall.

Increasing the sample size from 500 to 5,000 reduced disagreement by 0.19 percentage points and increased agreement with the limiting assignment by 0.16 percentage points. Increasing it further to 10,000 reduced disagreement and increased agreement with the limiting assignment by 0.03 percentage points each, while doubling computation time. HYphlow's default of 5,000 ancestral state samples was therefore retained.
